# Avirulence depletion assay: combining *R* gene-mediated selection with bulk sequencing for rapid avirulence gene identification in wheat powdery mildew

**DOI:** 10.1101/2024.07.10.602895

**Authors:** Lukas Kunz, Jigisha Jigisha, Fabrizio Menardo, Alexandros G. Sotiropoulos, Helen Zbinden, Shenghao Zou, Dingzhong Tang, Ralph Hückelhoven, Beat Keller, Marion C. Müller

## Abstract

Wheat production is threatened by multiple fungal pathogens, such as the wheat powdery mildew fungus (*Blumeria graminis* f. sp. *tritici*, *Bgt*). Wheat resistance breeding frequently relies on the use of resistance (*R*) genes that encode diverse immune receptors which detect specific avirulence (*AVR*) effectors and subsequently induce an immune response. While *R* gene cloning has accelerated recently, *AVR* identification in many pathogens including *Bgt* lags behind, preventing pathogen-informed deployment of resistance sources. Here we describe a new “avirulence depletion (AD) assay” for rapid identification of *AVR* genes in *Bgt*. This assay relies on the selection of a segregating, haploid F1 progeny population on a resistant host, followed by bulk sequencing, thereby allowing rapid avirulence candidate gene identification with high mapping resolution. In a proof-of-concept experiment we mapped the *AVR* component of the wheat immune receptor *Pm3a* to a 25kb genomic interval in *Bgt* harboring a single effector, the previously described *AvrPm3^a2/f2^*. Subsequently, we applied the AD assay to map the unknown *AVR* effector recognized by the Pm60 immune receptor. We show that *AvrPm60* is encoded by three tandemly arrayed, nearly identical effector genes that trigger an immune response upon co-expression with *Pm60* and its alleles *Pm60a* and *Pm60b*. We furthermore provide evidence that *Pm60* outperforms *Pm60a* and *Pm60b* through more efficient recognition of *AvrPm60* effectors, suggesting it should be prioritized for wheat breeding. Finally, we show that virulence towards *Pm60* is caused by simultaneous deletion of all *AvrPm60* gene paralogs and that isolates lacking *AvrPm60* are especially prevalent in the US thereby limiting the potential of *Pm60* in this region. The AD assay is a powerful new tool for rapid and inexpensive *AVR* identification in *Bgt* with the potential to contribute to pathogen-informed breeding decisions for the use of novel *R* genes and regionally tailored gene deployment.

## Introduction

Global wheat production is threatened by numerous pathogenic organisms with fungal diseases alone resulting in 15-20% yield losses annually (Figueroa *et al*., 2018; Savary *et al*., 2019). Sustainable wheat production therefore relies on extensive breeding efforts to identify new genetic resistance sources, including specific resistance (*R*) genes, and to introduce them into high yielding cultivars. Many *R* genes encode intracellular nucleotide-binding leucine-rich repeat (NLR) immune receptors that recognize the presence of pathogen effectors, so called avirulence (AVR) proteins, and subsequently induce an immune response. AVR recognition by NLRs often results in a hypersensitive response (HR) that includes localized cell death and thereby limits pathogen proliferation (Dodds & Rathjen, 2010; Ngou *et al*., 2022). Evasion of NLR recognition by mutation or loss of *AVR* genes is often observed in fast evolving plant pathogens and thereby significantly limits the durability of *R* genes deployed in agricultural settings (Brown, 2015). Several recent studies have highlighted the need for *AVR* identification and analysis of *AVR* diversity within pathogen populations in order to predict *R* gene durability and effective deployment in agricultural settings (Hafeez *et al*., 2021; Müller *et al*., 2022; Minter & Saunders, 2023).

While *R* gene identification and cloning in wheat and other staple crops has significantly sped up in recent years due to technological advances and vastly improved genomic resources (Running & Faris, 2024), identification of *AVR* genes lags behind for many pathogens and new, more efficient methods for *AVR* gene identification are urgently needed (Minter & Saunders, 2023).

Wheat powdery mildew (*Blumeria graminis* f.sp. *tritici,* abbrev. *Bgt*) is an obligate biotrophic ascomycete fungus that exhibits a high level of host specificity and exclusively infects wheat (Kusch *et al*., 2024). Due to its short asexual life cycle it can cause explosive epidemics among susceptible wheat monocultures and result in considerable yield losses (Beest *et al*., 2008). Until today, 69 *R* genes with over 100 functional alleles against *Bgt* have been genetically defined in wheat (*Pm1* to *Pm69*) with many more *R* gene candidates awaiting official definition (McIntosh *et al*., 2019; Hafeez *et al*., 2021). However, only a fraction of all defined *Pm* genes have so far been molecularly isolated and cloned with many of them (*Pm1a*, *Pm2*, *Pm3a-t*, *Pm5e*, *Pm8*, *Pm17*, *Pm21*, *Pm41*, *Pm60* and *Pm69*) encoding classic NLR proteins (Yahiaoui *et al*., 2004; Hurni *et al*., 2013; Sánchez-Martín *et al*., 2016; Singh *et al*., 2018; Zou *et al*., 2018; Hewitt *et al*., 2020; Li *et al*., 2020; Xie *et al*., 2020; Li *et al*., 2024). While some of the cloned *Pm* genes provide resistance against a narrow range of *Bgt* isolates or have been largely overcome by the pathogen population during agricultural deployment, other *Pm* genes such as *Pm60* provide resistance against most *Bgt* isolates and therefore represent valuable *R* gene candidates for future breeding efforts (Zou *et al*., 2018; Zhao *et al*., 2020).

*Pm60* has been identified and cloned from the wheat progenitor *Triticum urartu* and was found to encode an NLR (Zou *et al*., 2018). A recent study investigating the genetic diversity of the *Pm60* locus in *T. urartu* has furthermore revealed two additional *Pm60* alleles, *Pm60a* and *Pm60b*, that provide resistance against *Bgt* (Zhao *et al*., 2020). Interestingly, *Pm60a* is defined by a 240bp deletion, encompassing two entire LRR repeats. By contrast, *Pm60b* carries a 240bp insertion and therefore 2 additional LRR repeats when compared with *Pm60* (Zhao *et al*., 2020). Based on the resistance spectrum found in *Pm60*, *Pm60a* and *Pm60b* containing lines, it was hypothesized that the three alleles might recognize similar *AVR* effector targets albeit likely with differences in recognition strength (Zhao *et al*., 2020). However, the identity of the corresponding *AVR* effector *AvrPm60* from *Bgt* is so far unknown, which hampered detailed investigations of *Pm60* allele specificity and predictions of their exact potential for wheat breeding.

In the last decade, multiple *AVR* genes have been identified and cloned from *Bgt*: *AvrPm1a_1*, *AvrPm1a_2*, *AvrPm2*, *AvrPm3^a2/f2^*, *AvrPm3^b2/c2^*, *AvrPm3^d3^*, *AvrPm8*, *AvrPm17* were shown to be recognized by the NLRs *Pm1a*, *Pm2*, *Pm3a/Pm3f*, *Pm3b/Pm3c*, *Pm3d*, *Pm8* and *Pm17,* respectively (Bourras *et al*., 2015; Praz *et al*., 2017; Bourras *et al*., 2019; Hewitt *et al*., 2020; Müller *et al*., 2022; Kloppe *et al*., 2023; Kunz *et al*., 2023). Interestingly, all *Bgt AVR* effector genes known to date encode small, secreted effector proteins with a size of approximately 120 amino acids and are predicted to exhibit an RNAse-like structure (Bauer *et al*., 2021; Cao *et al*., 2023).

The identification and functional validation of *AVR/R* gene pairs combined with analyses of *AVR* diversity within the worldwide *Bgt* population has proven crucial to advance the understanding of gain-of-virulence mechanisms in *Bgt* and consequently resistance gene breakdown in wheat agriculture. For instance, two recent studies on the quick breakdown of the *Pm8* and *Pm17* resistance genes, introgressed from rye, found evidence for ancient genetic variation in the corresponding *AVR* genes within the *Bgt* population, including virulence alleles that evade *R* gene recognition. Importantly, these ancient *AVR* gene variants precede the introgression of *Pm8* and *Pm17* from rye into the wheat gene pool, explaining their rapid breakdown shortly after agricultural deployment (Müller *et al*., 2022; Kunz *et al*., 2023). Such examples highlight the importance of parallel *AVR* and *R* gene identification in pathogen and host in order to predict durability of *R* genes and to allow prioritization of most promising gene candidates for wheat breeding or *R* gene stacking.

*AVR* gene identification in *Bgt* has relied on a variety of experimental strategies using genetic mapping, GWAS, effector screens and, most recently, UV mutagenesis to induce and identify gain-of-virulence mutations (Bourras *et al*., 2015; Bourras *et al*., 2019; Bernasconi *et al*., 2024). Due to its haploid genome, the fast generation time and the experimentally controllable asexual (i.e. clonal) and sexual reproduction, genetic mapping approaches have proven particularly powerful in *Bgt* and remain the most used tool for *AVR* gene identification. However, genetic mapping using biparental crosses traditionally involves time-consuming isolation of 100+ individual F1 progeny from sexually formed chasmothecia, their subsequent asexual propagation, genotyping and phenotyping thereby resulting in work- and cost-intensive projects with timelines of 1-2 years. Hence, there is a need for technological advances to speed up *AVR* identification in this important wheat pathogen.

In this study, we describe a new “*AVR* depletion assay” (AD assay) for rapid and low-cost *AVR* gene identification in *Bgt*. The assay preserves the many advantages of genetic mapping approaches but circumvents the time-consuming isolation and propagation of individual F1 progenies by combining the generation of sexual recombinant F1 populations with *R* gene-mediated selection and bulk sequencing. In a proof-of-concept experiment, we show that the AD assay can identify the previously described *AvrPm3^a2/f2^* with high precision (i.e. identifying a single candidate effector). Furthermore, we apply the new method to identify and functionally validate the previously unknown *AvrPm60*, recognized by the broadly acting *Pm60* resistance gene.

## Results

### The avirulence depletion assay allows mapping of *AvrPm3^a2/f2^* with high resolution

In this study, we aimed at developing an “avirulence depletion assay” (AD assay) as a new approach for the identification of avirulence factors in *Bgt*. The assay is based on principles of bi-parental mapping but without the need to establish, maintain, and genotype individuals of a mapping population (Figure 1). As such, the approach relies on crossing two parental *Bgt* isolates displaying opposite virulence phenotypes on an *R* gene-containing wheat line and subsequent regeneration of a sexual F1 mapping population. In contrast to classical bi-parental mapping approaches (map-based cloning, QTL mapping), in the AD pipeline, conidiospores of a mixed F1 progeny population are directly used to infect a wheat line that carries the resistance of interest thereby creating a strong and directional selection pressure that depletes the F1 progeny population from individuals carrying the corresponding *AVR* factors. In parallel, the initial F1 progeny population is used to infect a susceptible wheat genotype subsequently serving as an unselected control. Following bulk harvesting and sequencing of the surviving F1 progenies, single nucleotide polymorphisms (SNPs) are used to identify regions with a deviation of the 1:1 parental genotype ratio expected for positions unaffected by any selection. Finally, the identified candidate regions are inspected using genomics and transcriptomics datasets to identify candidate *AVR* genes (Figure 1).

**Figure 1:**
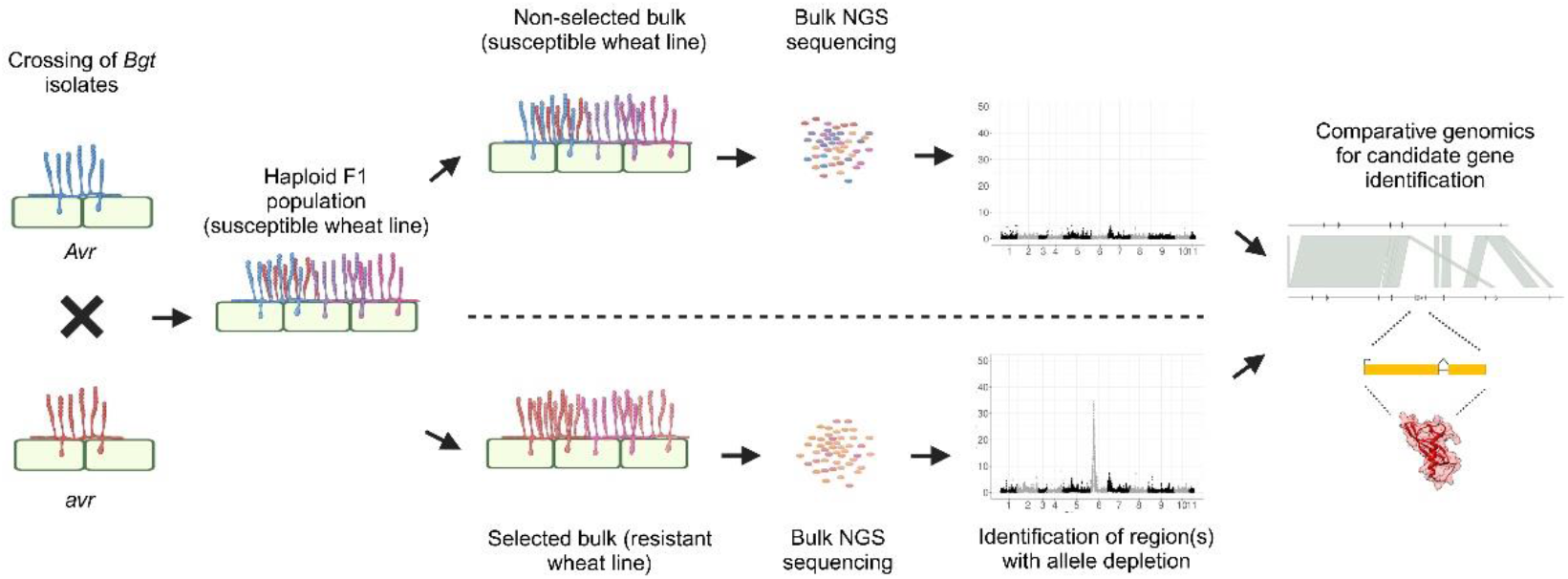
Schematic summary of the *AVR* depletion assay (AD assay) workflow. A bi-parental cross between *Bgt* isolates exhibiting opposite (a)virulence phenotypes results in a haploid F1 population which subsequently is selected on a resistant wheat line resulting in *AVR* depletion (selected bulk) or a susceptible control line (non- selected bulk). Bulk NGS sequencing and bioinformatic analyses are used to define genomic regions with *AVR* depletion and subsequently identify *AVR* candidate genes. Figure created with BioRender.com.

In a first experiment, we aimed to establish a proof-of-concept dataset for the AD-assay by using the well-characterized *Bgt* avirulence gene *AvrPm3^a2/f2^* located on Chr-06. *AvrPm3^a2/f2^*encodes an RNAse-like effector recognized by the wheat NLR allele *Pm3a* (Bourras *et al*., 2015). To test if the AD-assay can re-discover *AvrPm3^a2/f2^*, we crossed the Swiss isolate CHVD_042201 (*AvrPm3a*, MAT 1-2) with the Chinese isolate CHN_52_27 (*avrPm3a*, MAT 1-1) that display opposite phenotypes on the *Pm3a* containing near-isogenic line ‘Asosan/8*CC’ (Figure 2a). From this cross, we generated a mixed population of an estimated 1500 F1 progenies on the susceptible wheat line ‘Kanzler’, which is devoid of *R* genes against *Bgt*. In the next step, we used conidiospores of this multiplied F1 progeny population to infect either ‘Asosan/8*CC’, thereby creating a *Pm3a*-mediated selection pressure (*Pm3a*-selected bulk) or the susceptible wheat cultivar ‘Kanzler’ to create an unselected control bulk. Subsequently, we bulk-harvested conidiospores produced during the respective selection steps and subjected both bulks to DNA sequencing using Illumina paired-end reads to a coverage of approximately 120X.

**Figure 2:**
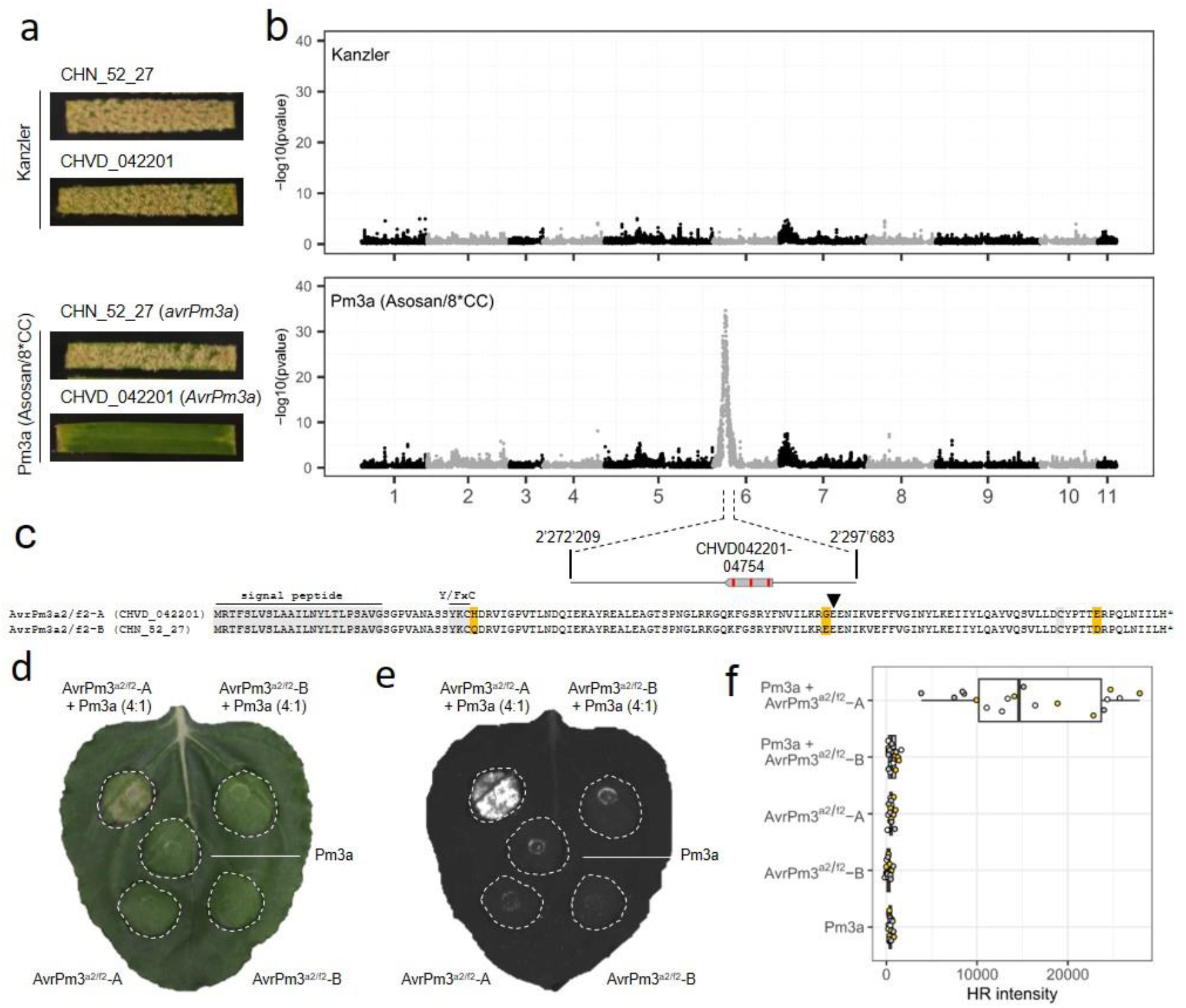
High-resolution mapping and functional validation of the previously described *AvrPm3^a2/f2^* in a proof- of-concept study using the newly developed AD assay. **(a)** Virulence phenotypes of the parental *Bgt* isolates CHN_52_27 and CHVD_042201 on the susceptible wheat line ‘Kanzler’ and the *Pm3a*-containing line ‘Asosan/8*CC’. **(b)** Statistical analysis of sequenced bulks selected on ‘Kanzler’ or ‘Asosan/8*CC’ to detect regions with deviations from an expected 1:1 parental genotype ratio. Individual datapoints represent –log10 transformed p-values of a G-test at individual SNPs marker positions. A single genomic region on Chr-06 exhibits deviations from an expected 1:1 parental genotype inheritance in the *Pm3a*-selected bulk. The mapped *AvrPm3a* target interval (≥95% reads from virulent parent) encompassing 25kb and the single effector gene *CHVD042201- 04754* (*AvrPm3^a2/f2^*) is depicted in a zoom-in (bottom). **(c)** Protein sequence alignment of AvrPm3^a2/f2^-A found in the avirulent parental isolate CHVD_042201 and AvrPm3^a2/f2^-B in the virulent parental isolate CHN_52_27. Amino acid polymorphisms are highlighted in yellow. The signal peptide, the conserved Y/FxC motif, the C-terminal cysteine and the conserved position of a small intron, all serving as hallmarks of RNAse-like effectors are highlighted in grey or by a small arrowhead (intron position). (d, e) *Agrobacterium*-mediated expression of AvrPm3^a2/f2^-A, AvrPm3^a2/f2^-B and Pm3a in *N. benthamiana*. Co-infiltrations (top) were performed with a 4 (effector) : 1 (NLR) ratio. Leaves were imaged using a camera (d) or the Fusion FX imager system (e) at four days post inoculation. The assay was performed with n=6 leaves and repeated a total of three times with similar results (total n=18 leaves). **(f)** Quantification of HR intensity in the *N. benthamiana* expression assay depicted in (e). Datapoints from three independent experiments with n=6 leaves are color coded (total n=18).

The *AvrPm3^a2/f2^* gene is located in an effector cluster consisting of 19 members of the E008 candidate effector family (Bourras *et al*., 2015). Due to its complexity, the locus was not fully resolved in the previously published *Bgt* reference genome assembly of strain CHE_96224, where it was found to contain several sequence gaps and collapsed regions for highly similar gene copies (Müller *et al*., 2019). Given the advances in long-read sequencing technologies, we saw the opportunity to generate an improved reference genome assembly that fully resolves the *AvrPm3^a2/f2^* locus and potentially other collapsed or incomplete regions in the published *Bgt* reference genome CHE_96224. To do so, we sequenced CHVD_042201, which served as an avirulent parent in our AD assay, with PacBio HiFi to a coverage of 100X and assembled the genome using the hifiasm assembler (see Supplementary Note 1 for details). This strategy resulted in a near-complete telomere-to-telomere assembly with fully resolved centromeres and only two remaining sequence gaps in the highly expanded and repetitive rRNA encoding region on Chr-09 and a previously identified array of tandem repeats on Chr-04. Importantly, the new genome assembly also fully resolved the *AvrPm3^a2/f2^* locus and multiple other previously collapsed regions and thus represents a significant improvement in terms of genome resolution and continuity compared to previous *Bgt* assemblies (Supplementary Table 1, Supplementary Note 1).

To analyse the bulked DNA sequencing data generated as part of the AD assay pipeline, we mapped the Illumina reads from the *Pm3a*-selected bulk, the non-selected control bulk, and the parental isolates CHVD_042201 and CHN_52_27 on the new genome assembly of CHVD_042201 and identified 198’027 high-quality SNPs between the two parental isolates serving as genetic markers in the subsequent analysis. In the sequenced bulks, we expected regions unaffected by any selection to show a 1:1 ratio of parental genotypes at any given marker. In contrast, in regions under selection, we expected ratios to significantly deviate from a 1:1 ratio, with the genotype of the avirulent parent CHVD_042201 being underrepresented (i.e. depleted). As expected, we did not identify any region with a strong deviation from a 1:1 ratio for the control bulk selected on the susceptible cultivar ‘Kanzler’ that is devoid of any *R* genes against *Bgt* (Figure 2b). In contrast, in the *Pm3a*-selected bulk, we found a single region located on Chr-06 that showed strong deviation from a 1:1 ratio towards the genotype from the virulent CHN_52_27 parental isolate (Figure 2b). The identified genomic segment overlapped with the previously identified E008 candidate effector cluster containing the *AvrPm3^a2/f2^* gene. Importantly, the region with the strongest depletion signal (i.e., ≥95% of reads from virulent parent) was constricted to 25kb and encompassed a single gene CHVD042201-04754 (Figure 2b). The identified gene is identical to the *AvrPm3^a2/f2^* variant *AvrPm3^a2/f2-A^-A*, which was previously shown to be recognized by *Pm3a* (Bourras *et al*., 2015; McNally *et al*., 2018). Based on resequencing data, we determined that the CHVD042201-04754 homolog in the *Pm3a-* virulent isolate CHN_52_27 encodes for a protein variant with three amino acid changes compared to CHVD_042201 (H36Q, G84E, E121D) (Figure 2c). This haplovariant was previously identified in isolates from China and Israel and was termed *AvrPm3^a2/f2^*-B and designated as a putative *AvrPm3^a2/f2^* gain-of-virulence allele (McNally *et al*., 2018). To confirm this finding, we expressed the *AvrPm3^a2/f2^* variants from the two parental isolates (without signal peptide) together with *Pm3a* in *Nicotiana benthamiana* using *Agrobacterium*-mediated transient transformation. Consistent with the results from the AD-assay, CHVD042201-04754 (*AvrPm3^a2/f2^-A*) elicited a *Pm3a*-dependent hypersensitive response (HR), whereas the *AvrPm3^a2/f2^-B* from the virulent parent CHN_52_27 did not, thereby confirming *AvrPm3^a2/f2^-B* to be a virulence allele (Figure 2d-f). Based on these findings, we concluded that the avirulence allele depletion observed in the *Pm3a*-selected bulk is a direct consequence of differential recognition of AvrPm3^a2/f2^ variants found in the two parental isolates. In conclusion, our proof-of-concept datasets showed that the AD-assay is a powerful new tool to identify avirulence factors in *Blumeria* down to single gene resolution.

For this proof-of-concept study of the AD assay, we relied on the high-quality genome assembly of CHVD_042201, which represented the avirulent parental isolate in our genetic cross. Even though the availability of a complete genome sequence of the avirulent parent is likely ideal to identify candidates genes, we wanted to test whether the AD assay succeeds in *AVR* identification also with an alternative reference genome. We therefore tested our AD pipeline using the previously published reference genome assembly of isolate CHE_96224 (Bgt_genome_v3_16) (Müller *et al.,* 2019). Similar to the above-described analysis based on CHVD_042201, we found no deviations from the expected 1:1 parental genotype ratio in the unselected F1 bulks but a single region with strong avirulence depletion signal on Chr-06 in the *Pm3a*-selected bulk (Supplementary Figure S1, Supplementary Table 2, Supplementary Note 2). The identified genomic region in the CHE_96224 overlapped with the identified *AvrPm3^a2/f2^*locus in CHVD_042201, thereby confirming that the AD approach succeeds in identifying *AvrPm3^a2/f2^* also with an alternative reference genome assembly (see Supplementary Note 2 for details).

### The AvrPm60 effector is encoded by three tandem duplicated genes in the *Pm60*-avirulent isolate CHVD_042201

Next, we aimed to use the AD assay to identify the so far unknown *AvrPm60*, the avirulence factor corresponding to the NLR *Pm60 (Zou et al., 2018)*. The parental *Bgt* isolates CHVD_042201 and CHN_52_27 displayed opposite virulence phenotypes on ‘Kn199 *Pm60’* (Figure 3a), a transgenic line expressing the *Pm60* gene in the susceptible background ‘Kn199’ (Zou *et al*., 2018). We therefore used the ‘Kn199 *Pm60’* transgenic line to apply a *Pm60*- mediated selection to the CHVD_042201 x CHN_52_27 F1 progeny population described above. Again, the AD- assay identified a single genomic region in which the *Pm60*-selected bulk showed a strong deviation from the 1:1 parental genotype ratio, with a depletion of the avirulent CHVD_042201 parental genotype. Strikingly, the *AvrPm60* candidate locus partially overlapped with the above-described *AvrPm3^a2/f2^* locus on Chr-06 (Figure 3b). A parallel analysis of the *Pm60*-selected bulk based on the alternative CHE_96224 reference genome assembly identified a single genomic region with strong depletion of the avirulent genotype on Chr-06 which overlapped with the identified region in CHVD_042201, again highlighting the independence of the AD assay from individual reference genomes (Supplementary Figure S1, Supplementary Table 2, Supplementary Note 2).

**Figure 3:**
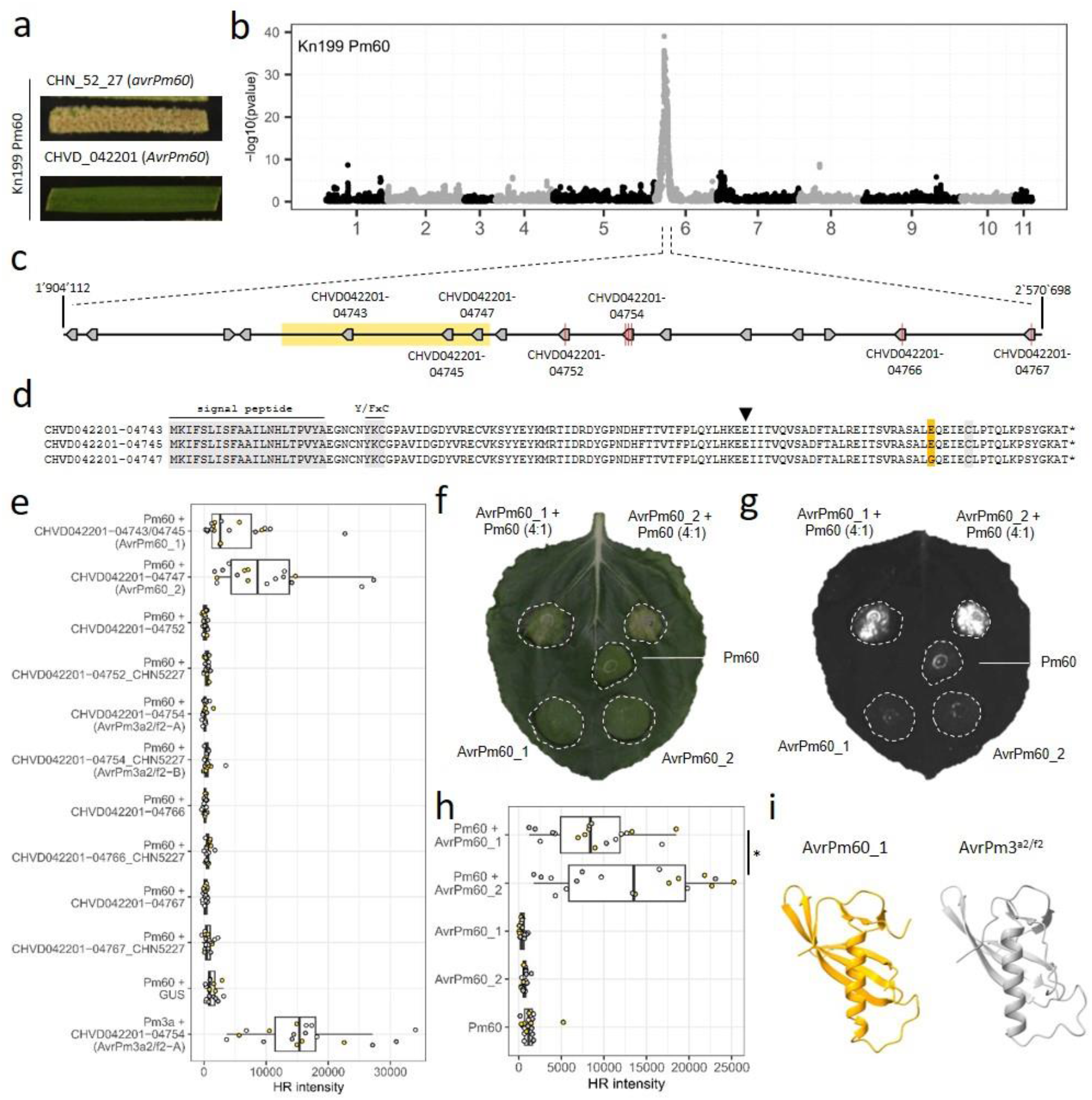
Mapping and functional validation of *AvrPm60_1* and *AvrPm60_2* using the AD assay. **(a)** Virulence phenotype of the parental *Bgt* isolates CHN_52_27 and CHVD_042201 on the transgenic wheat line ‘Kn199 Pm60’. **(b)** Statistical analysis of sequenced bulks selected on ‘Kn199 *Pm60*’ to detect regions with deviations from an expected 1:1 parental genotype ratio. Individual datapoints represent –log10 transformed p-values of a G-test at individual SNPs marker positions. A single genomic region on Chr-06 exhibits deviations from an expected 1:1 parental genotype inheritance in the *Pm60*-selected bulk. **(c)** Schematic zoom-in of the mapped *AvrPm60* target interval (≥95% reads from virulent parent) encompassing 667 kb and 16 candidate effector genes. Polymorphic candidate effectors are individually labelled, and non-synonymous SNPs indicated by a red line. The genomic region encompassing *CHVD042201-04743, CHVD042201-04745* and *CHVD042201-04747,* which was found to be deleted in the virulent parental isolate CHN_52_27, is highlighted in yellow. **(d)** Protein sequence alignment of the identical effectors CHVD042201-04743/CHVD042201-04745 and CHVD042201-04747 (E103G). The E103G polymorphism is highlighted in yellow. The signal peptide, the conserved Y/FxC motif, the C- terminal cysteine and the conserved position of a small intron, all serving as hallmarks of RNAse-like effectors are highlighted in grey or by a small arrowhead (intron position). **(e)** Co-expression of polymorphic effector candidates found within the mapped *AvrPm60* locus together with Pm60 using *Agrobacterium-*mediated expression in *N. benthamiana*. Co-expression of GUS + Pm60 and AvrPm3^a2/f2^-A + Pm3a were used as negative and positive controls, respectively. Co-infiltrations were performed with a 4 (effector) : 1 (NLR) ratio and imaged with a Fusion FX imager system 4 days post inoculation. HR intensity was quantified from three independent experiments with n=6 leaves (color-coded datapoints, total n=18). **(f, g)** *Agrobacterium*-mediated expression of AvrPm60_1, AvrPm60_2 and Pm60 in in *N. benthamiana*. Co-infiltrations (top) were performed with a 4 (effector) : 1 (NLR) ratio. Leaves were imaged using a camera (f) or the Fusion FX imager system (g) at four days post inoculation. The assay was performed with n=6 leaves and repeated a total of three times with similar results (total n=18 leaves). **(h)** Quantification of HR intensity in the *N. benthamiana* expression assay depicted in (g). Datapoints from three independent experiments with n=6 leaves are color coded (total n=18). * indicates statistical difference according to a Wilcoxon signed rank test (p<0.05). **(i)** Predicted three dimensional structures of AvrPm60_1 (yellow) and AvrPm3^a2/f2^ (grey) according to Alphafold 3 structural modelling. Both effector proteins are predicted to exhibit an RNAse-like structure.

The region with the strongest signal of avirulence allele depletion (i.e. ≥95% of the reads originated from the virulent parent CHN_52_27) in the *Pm60*-selected bulk encompassed a region of 667kb in CHVD_042201 containing 16 annotated, high-quality genes, all belonging to the E008 candidate effector family (Figure 3c). We inspected all genes in the locus with re-sequencing data from CHVD_042201 and CHN_52_27 and found that only seven of the 16 candidate effectors, including the above-described *AvrPm3^a2/f2^* gene (CHVD042201-04754) exhibited polymorphisms between the parental isolates and therefore represented good *AvrPm60* candidate genes (Figure 3c). Among the seven polymorphic genes, only four carried SNPs resulting in amino acid polymorphisms between the virulent and avirulent parents. For the remaining three genes (*CHVD042201-04743, CHVD042201-04745, and CHVD042201-04747*), we did not detect any alignment of sequencing reads from CHN_52_27, indicating that these three genes are deleted in the virulent parent (Figure 3c). Interestingly, *CHVD042201-04743, CHVD042201-04745*, and *CHVD042201-04747* constitute tandem duplicates of the same effector gene in the avirulent isolate CHVD_042201. The resulting effector proteins CHVD042201-04743 and CHVD042201-04745 are identical, whereas CHVD042201-04747 differs by a single amino acid change (E103G) (Figure 3d). To functionally validate *AvrPm60*, we co-expressed the seven polymorphic candidate genes found within the mapped locus together with *Pm60* using transient overexpression in *N. benthamiana*. Both the CHVD042201-04743/CHVD042201-04745 and CHVD042201-04747 effectors triggered a *Pm60*-dependent HR response, whereas none of the other candidates or a GUS negative control resulted in cell death (Figure 3e-h). Hence, we concluded that three genes *CHVD042201-04743, CHVD042201-04745*, and *CHVD042201-04747* represent *AvrPm60* by encoding two nearly identical effector proteins that we designated as AvrPm60_1 (CHVD042201-04743/CHVD042201-04745) and AvrPm60_2 (CHVD042201-04747). Interestingly, the AvrPm60_2 variant carrying the amino acid polymorphism E103G elicited a slightly stronger HR response compared to AvrPm60_1 (Figure 3e-h).

### AvrPm60_1 and AvrPm60_2 belong to the RNAse-like effector superfamily

The identified AvrPm60 effectors are part of the large effector family E008 with over 40 members, including the AvrPm3^a2/f2^ effector (Müller *et al*., 2019). Like other member of the E008 family, the two AvrPm60 are small proteins of 121 amino acids in size, with the first 23 amino acids constituting a predicted signal peptide (Figure 3d). Similar to all previously identified AVRs in *Bgt*, the AvrPm60 proteins contain a Y/FxC motif and a conserved C-terminal cysteine, two features that were defined as hallmarks of RNAse-like effectors which comprise more than half of all effector proteins found in the *Blumeria* genus (Cao *et al*., 2023; Seong & Krasileva, 2023). Indeed, structural modelling using Alphafold 3 (Abramson *et al*., 2024) predicted that the AvrPm60 proteins exhibit an RNAse-like structure and are structurally similar to the E008 family member AvrPm3^a2/f2^ (Figure 3i).

Multiple previously identified RNAse-like AVRs in *Bgt* were found to exhibit very high expression levels during early phases of infection (Bourras *et al*., 2015; Praz *et al*., 2017; Bourras *et al*., 2019; Kunz *et al*., 2023). Similarly, analysis of RNA sequencing data from five *Bgt* isolates at two days post inoculation (2dpi) showed that the *AvrPm60* genes are consistently among the top 5% of the highest expressed genes in each isolate (Supplementary Figure S2). In summary, the AvrPm60 effectors are bona-fide members of the RNAse-like effector class in *Blumeria*.

### Gain-of-virulence through deletion of *AvrPm60* genes is rare within the worldwide *Bgt* population but widespread in the US

The *Pm60*-virulent isolate CHN_52_27 used in the AD assay evades *Pm60*-mediated resistance due to a large-scale deletion encompassing all three *AvrPm60* copies found in CHVD_042201 (Figure 3c). We therefore aimed to investigate the frequency and distribution of this striking gain-of-virulence mechanism within the global *Bgt* population. To estimate the number of *AvrPm60* gene copies, we used a previously described *in-silico* approach using publicly available resequencing data from 382 *Bgt* isolates (Menardo *et al*., 2016; Praz *et al*., 2017; Müller *et al*., 2019; Müller *et al*., 2022; Sotiropoulos *et al*., 2022; Kloppe *et al*., 2023). This approach uses normalized read coverage as a proxy for the number of gene copies in each isolate. As a control we used the *GAPDH* gene which was previously shown to be present as a single copy gene, and *AvrPm3^a2/f2^* found to occur in 1-4 copies in *Bgt* isolates (Müller *et al*., 2019). As expected, the coverage analysis indicated that GAPDH occurs as a single copy gene in all 382 analysed isolates, whereas the analysis found signs of copy number variations for both the *AvrPm3^a2/f2^* and the *AvrPm60* gene in this worldwide *Bgt* panel (Figure 4a, Supplementary Figure S3). Consistent with the literature, the coverage analysis indicates that all isolates carry at least one copy of the *AvrPm3^a2/f2^*, although higher-order duplications with two or more copies are readily observed (Supplementary Figure S3), (McNally *et al*., 2018; Müller *et al*., 2019; Müller *et al*., 2021). In contrast, we estimated that the majority of *Bgt* isolates contain two *AvrPm60* copies (Figure 4a). However, we detected sizeable additional copy number variation in the *Bgt* diversity panel with some isolates containing a single *AvrPm60* gene and isolates with three or more *AvrPm60* copies. Consistent with the broad functionality of *Pm60* described in the literature (Zou *et al*., 2018), only a minority of isolates (13/382), including the Chinese isolate CHN_52_27, were devoid of any *AvrPm60* copies as indicated by the absence of any sequencing coverage in our analysis (Figure 4a, Table 1). Interestingly, among the 13 isolates lacking *AvrPm60* only one additional isolate originated from China (2 out of 63 Chinese isolates), while the remaining isolates all originated from the US, where 19% of investigated isolates carried the *AvrPm60* deletion (Table 1). To experimentally confirm the deletion of *AvrPm60* gene copies in specific isolates from China and the US, we designed *AvrPm60*-specific PCR primers based on conserved flanking sequences of all three *AvrPm60* genes in the reference isolate CHVD_042201. Using these primers we successfully amplified *AvrPm60* from genomic DNA of CHVD_042201 and six additional *Bgt* isolates with diverse geographic origin (Switzerland, UK, China, Japan, Argentina, US) for which our coverage analysis estimated between one and four *AvrPm60* copies in the genome. In contrast, PCR amplification failed from CHN_52_27 and three isolates from the US with a predicted complete *AvrPm60* deletion, thereby confirming the results of the coverage analysis (Supplementary Figure S4). We then subjected the same 11 *Bgt* isolates to virulence phenotyping on the transgenic wheat line ‘Kn199 + Pm60’ and the susceptible control ‘Kn199’. Importantly, all isolates with at least one *AvrPm60* gene copy in the genome exhibited an avirulent phenotype on the *Pm60* transgenic line similar to CHVD_042201, whereas all tested isolates with *AvrPm60* gene deletions exhibited full virulence on *Pm60* wheat, comparable to the virulent CHN_52_27 isolate (Figure 4b). Hence, we conclude that deletion of *AvrPm60* genes represents a gain-of-virulence mechanism that allows *Bgt* to overcome *Pm60*- mediated resistance and that such *AvrPm60* gene deletions are particularly prevalent in the US, likely limiting *Pm60* efficacy in this geographic area.

**Figure 4:**
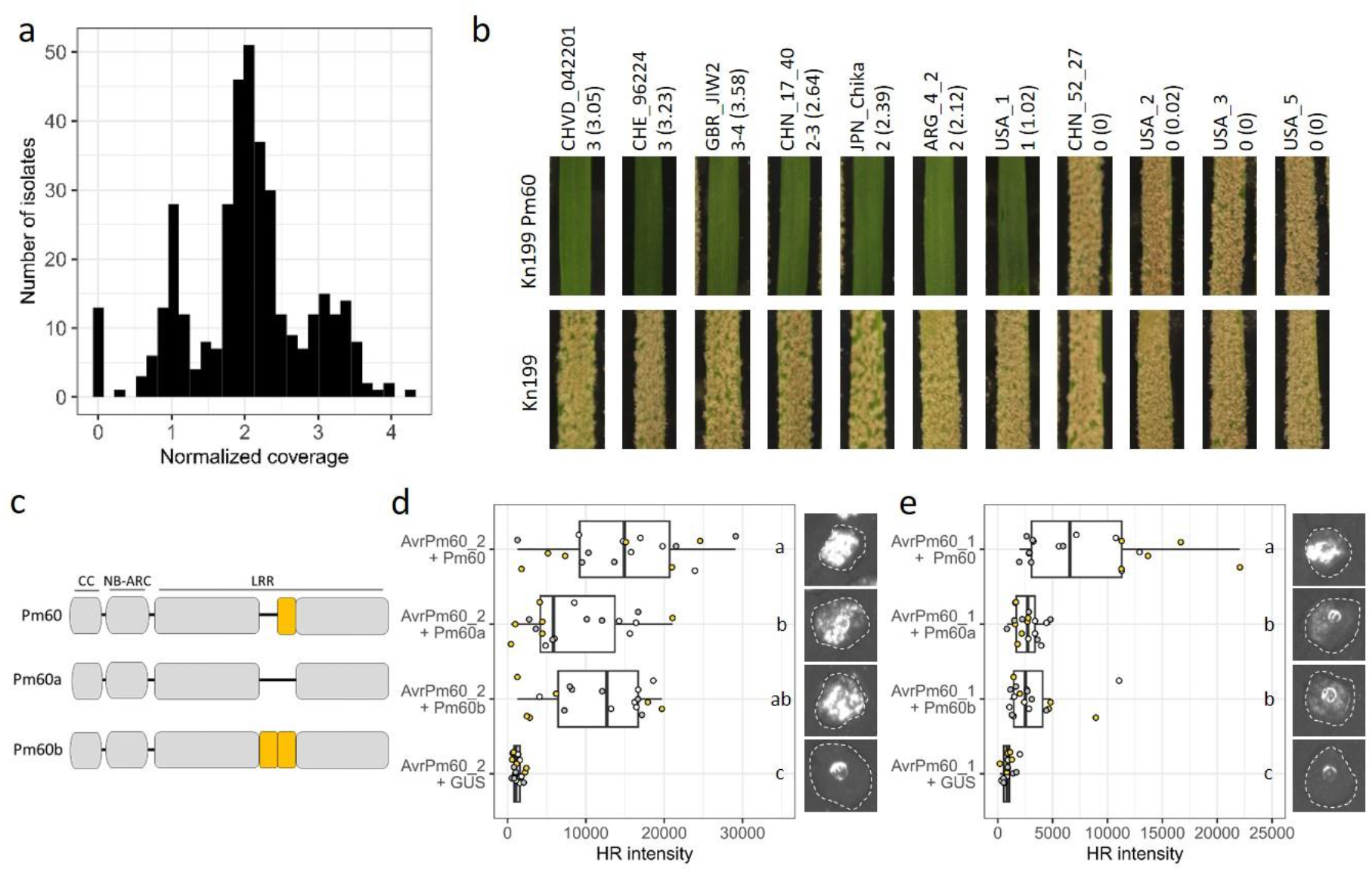
Functional characterization of *AvrPm60* copy number variation and *Pm60* allelic variants. **(a)** Copy-number estimation of *AvrPm60* gene copies in a worldwide diversity panel of 382 publicly available sequenced *Bgt* isolates. Genomic coverage of Illumina reads aligning to *AvrPm60* genes copies were normalized to the coverage of all genes in the genome. **(b)** Virulence phenotypes of 11 *Bgt* isolates with varying copy numbers of *AvrPm60* genes on the transgenic wheat line ‘Kn199 Pm60’ and ‘Kn199’, serving as a susceptible control. The estimated number of *AvrPm60* gene copies according to sequencing coverage analysis are indicated next to the isolate name with normalized sequencing coverage indicated in brackets. The isolates CHVD_042201 and CHN_52_27 used for initial *AvrPm60* identification are shown as comparison. **(c)** Schematic representation of the three tested *Pm60* alleles originating from *Triticum urartu*. Yellow boxes indicate a pair of LRR repeats that show copy number variations between the tested Pm60 alleles. **(b,c)** *Agrobacterium*-mediated expression of AvrPm60_1 (c), AvrPm60_2 (b) with Pm60, Pm60a and Pm60b in *N. benthamiana*. Co-infiltrations were performed with a 4 (effector) : 1 (NLR) ratio. Leaves were imaged at 4dpi using the Fusion FX imager system. The assay was performed with n=6 leaves and repeated a total of three times with similar results (total n=18 leaves). Boxplots represent quantification of HR intensity in the *N. benthamiana* expression assay. Datapoints from the three independent experiments are color coded. Different letters next to the boxplot represent statistical differences according to a pairwise Wilcoxon rank sum exact test (p<0.05).

**Table 1:** Estimated copy number of *AvrPm60* genes in *Bgt* isolates originating from different geographic regions.

| origin | # isolates | deletion | 1 copy | 2 copies | >2 copies | unambiguous |
| --- | --- | --- | --- | --- | --- | --- |
| Argentina | 8 | - | 3 | 5 | - | - |
| China | 63 | 2 | 2 | 30 | 24 | 5 |
| Egypt | 48 | - | - | 33 | 6 | 9 |
| Europe | 30 | - | 5 | 5 | 20 | - |
| Iran | 10 | - | - | 7 | 3 | - |
| Israel | 31 | - | 2 | 16 | 11 | 2 |
| Japan | 10 | - | 4 | 5 | - | 1 |
| Kazakhstan | 3 | - | - | 1 | 2 | 0 |
| Russia | 38 | - | 4 | 23 | 7 | 4 |
| Turkey | 83 | - | 5 | 63 | 10 | 5 |
| USA | 58 | 11 | 38 | 3 | - | 6 |
| Total | 382 | 13 | 63 | 191 | 83 | 32 |

### *Pm60* and its allelic variants *Pm60a* and *Pm60b* recognize *AvrPm60_1* and *AvrPm60_2* with varying efficacy

The *Pm60* resistance gene was originally isolated from *T. urartu*, the progenitor of the A genome of hexaploid wheat. Allele mining in *T. urartu* identified two additional functional alleles of *Pm60*, termed *Pm60a* and *Pm60b*, that provide resistance against *Bgt* (Zou *et al*., 2018; Zou *et al*., 2022). *Pm60a* and *Pm60b* differ from *Pm60* through the deletion or tandem duplication of two LRR repeats, respectively (Figure 4c) (Zou *et al*., 2018). In the literature, the *Bgt* recognition spectra of *Pm60a*, *Pm60b* and *Pm60* are described as largely overlapping thus prompting us to hypothesize that the three *Pm60* alleles recognize the same avirulence component in *Bgt*. To test this hypothesis, we co-expressed the three *Pm60* alleles with the *AvrPm60_1 and AvrPm60_2* in *N. benthamiana.* Both *AvrPm60* effector variants induced a hypersensitive response upon co-expression with *Pm60a, Pm60b* and *Pm60* but not with a *uidA* (GUS) negative control (Figure 4d,e) thus showing that the three *Pm60* alleles recognize the same avirulence components in *Bgt*. Although both *AvrPm60* variants are recognized by all three *Pm60* alleles, we observed significant differences in the strength of the hypersensitive response depending on the specific *Avr*/*R* combination. For the stronger *AvrPm60_2* variant, the *Pm60a* allele showed a marked reduction in the strength of HR when compared to *Pm60*. In contrast, recognition of the weaker *AvrPm60_1* variant resulted in reduced HR output for both *Pm60a* and *Pm60b* alleles when compared to *Pm60* (Figure 4d,e). In summary, our findings indicate that the three *Pm60* alleles, originating from *T. urartu*, recognize the same avirulence effectors AvrPm60_1 and AvrPm60_2, albeit with differences in the strength of the elicited HR. This indicates that the copy number variation in the LRR repeats does not influence the recognition specificity of the Pm60 alleles, but it might play a role in modulating the amplitude of the HR upon effector recognition.

## Discussion

Breeding genetically resistant cultivars is a cornerstone of sustainable agricultural production, with global efforts focusing on the identification of new resistance sources and their efficient deployment against fast evolving pathogenic organisms (Huang *et al*., 2023). In wheat, more than 460 resistance gene loci against various pathogens are genetically defined but only a small fraction of the underlying *R* genes have been cloned to date (McIntosh *et al*., 2019; Hafeez *et al*., 2021). Owing to technological advances and significantly improved genomic resources, the speed of *R* gene cloning in wheat has increased tremendously in recent years and it is expected that the majority of defined *R* genes will be molecularly identified within the next decade (Sánchez-Martín & Keller, 2021; Wulff & Krattinger, 2022). However, the question remains how these resistance sources can be deployed in modern agricultural systems to durably withstand continually evolving pathogen populations. In a widely acclaimed perspective paper, Hafeez and colleagues in 2021 called for a concerted, international effort to generate a wheat *R* gene atlas to tackle these problems (Hafeez *et al*., 2021). In the outlined vision, *R* gene identification in wheat must be accompanied by *AVR* identification and subsequent analysis of *AVR* diversity in corresponding pathogen populations in order to guide future breeding decisions and allowing for efficient, regionally tailored, *R* gene deployment. The authors argued that for the success of such an endeavor it is crucial to improve our current limited knowledge about *AVR* factors in wheat pathogens and in particular develop methods to speed up *AVR* gene identification in order to keep up with the improved pace of *R* gene cloning in the host (Hafeez *et al*., 2021).

In contrast to other wheat pathogens where few or no *AVR* factors have been identified to date, the field of *AVR* gene identification in *Bgt* is relatively advanced. Using various methods including classic genetic mapping, GWAS, mutagenesis and effector screens, a total of 8 *AVR* effectors have been cloned and molecularly characterized to date (Bourras *et al*., 2015; Praz *et al*., 2017; Bourras *et al*., 2019; Hewitt *et al*., 2020; Müller *et al*., 2022; Kloppe *et al*., 2023; Kunz *et al*., 2023; Bernasconi *et al*., 2024). Due to the haploid genome and experimentally accessible sexual reproduction of *Bgt*, classic genetic mapping has proven to be a powerful tool. For example, genetic mapping approaches can outperform GWAS in resolving genetically complex traits with multiple components or when components occur at low frequency in the *Bgt* population. Furthermore, in contrast to effector screening approaches, genetic mapping does not solely rely on HR as the primary immune outcome. However, the requirement to isolate, phenotype and genotype individual F1 progenies represents a major obstacle in genetic mapping experiments, leading to work- and cost-intensive projects in the range of 1-2 years. The newly developed AD assay described here leverages the many strengths of genetic mapping in *Bgt* while eliminating the work-intensive characterization of individual F1 progenies. It thereby shortens project timelines to approximately 4 months and reduces associated sequencing efforts to a selected and an unselected F1 pool resulting in much lower costs. Avoiding the most work-intensive steps in mapping projects furthermore allows *AVR* identification to be parallelized by either selecting the same progeny population on multiple *R* gene containing lines, as done in this study on *Pm3a* and *Pm60*, or even by simultaneously generating multiple progeny populations arising from different genetic crosses that segregate for different *AVR* factors.

Our proof-of-concept experiment aiming at the mapping of *AvrPm3^a2/f2^* identified a narrowly defined genomic region of 25kb harboring a single candidate effector gene (i.e. *AvrPm3^a2/f2^*), exemplifying the mapping power and resolution of the AD assay (Figure 2b). In the case of *AvrPm60* the mapped interval was however significantly bigger and encompassed a region of 667 kb with 16 effectors in total (7 polymorphic), indicating that mapping resolution varies depending on the mapped factor (Figure 3c). We hypothesize that these differences arise in part from the fact that *AvrPm60* is represented by three genes spread over 130 kb of sequence. The mapping resolution is likely also influenced by the rate of recombination within the mapped *AVR* locus, which was shown to differ throughout the *Bgt* genome (Müller *et al*., 2019) and the strength of selection exerted by the resistant wheat line used to generate the selected F1 bulks.

Interestingly, we observed near complete selection for the virulence allele (i.e. ≥95% of sequenced reads) in *Pm3a*- and *Pm60*-selected pools after a single asexual reproduction cycle likely due to the strong resistance effect exerted by both *R* genes. We are however optimistic that the AD assay could also be used to map *AVR* factors corresponding to *R* genes with weaker resistance effects, such as *Pm8* which was shown to result in partial resistance against isolates carrying *AvrPm8* under endogenous *Pm8* expression levels and only resulted in complete resistance in transgenic overexpression lines (Kunz *et al*., 2023). In such cases, the deviation from a 1:1 inheritance ratio in selected progeny pools might be less drastic, resulting in larger mapped intervals and consequentially a higher number of candidate genes. The situation could however be alleviated by extending the number of asexual cycles under selection or, if available, by the use of transgenic overexpression lines with stronger or complete resistance.

Several recent studies have shown that the genetic control of avirulence phenotypes in *Bgt* can be complex and can involve multiple genetic loci (Bourras *et al*., 2015; Hewitt *et al*., 2020; Müller *et al*., 2022; Kloppe *et al*., 2023). In the case of a rye introgression carrying *Pm17* and a genetically linked unknown second resistance gene, a QTL mapping approach using ∼120 F1 progenies successfully mapped both corresponding *AVR* loci in *Bgt* (Müller *et al*., 2022). However, in another study the complex avirulence landscape for several *Pm3* alleles was only partially resolved in a biparental mapping population of ∼140 F1 progenies (Bourras *et al*., 2015). It will be interesting to see whether the newly developed AD assay, due to the high number of processed progenies (∼1500 in this study), could improve mapping resolution in these complex cases that include multiple *AVR* loci. We hypothesize that the resolution achieved in the AD assay could be further improved by extending the F1 progeny population and in particular by sequencing bulks at significantly higher coverages (∼120X in this study), thereby capturing a more detailed picture of the recombination landscape within the F1 progeny population.

In conclusion, we argue that the AD assay is a powerful new tool for *AVR* identification in *Bgt* that will significantly speed up *AVR* cloning in this important wheat pathogen in the future. Furthermore, we hypothesize that the AD assay could be adapted to map avirulence components in other pathosystems, where the sexual reproduction cycle of the pathogen is experimentally accessible and genetic resources of the host include clearly defined *R* genes that allow for efficient selection of progeny pools.

Our knowledge about AVR effectors in cereal powdery mildews has improved significantly in recent years. With a total of 8 known and functionally validated *AVRs* in the wheat infecting *Bgt* and another 6 *AVRs* from barley powdery mildew (*Blumeria hordei*), common principles of AVR effectors in this group of cereal pathogens can be defined (Bourras *et al*., 2015; Lu *et al*., 2016; Praz *et al*., 2017; Bourras *et al*., 2019; Saur *et al*., 2019; Hewitt *et al*., 2020; Bauer *et al*., 2021; Müller *et al*., 2022; Kloppe *et al*., 2023; Kunz *et al*., 2023). Interestingly, all 14 AVRs belong to an effector superfamily of structurally similar proteins exhibiting an RNAse-like fold, but lacking RNAse activity (Bauer *et al*., 2021; Cao *et al*., 2023). Apart from the presence of a signal peptide, these proteins share several characteristics such as a N-terminal Y/FxC motif and a conserved C-terminal cysteine involved in disulfide bridge formation and hence stability of the RNAse fold (Pennington *et al*., 2019; Cao *et al*., 2023). Furthermore, RNAse-like effector genes share a single small intron at a conserved position indicating they evolved through diversification from the same ancestral gene, although they often share little amino acid identity (Pedersen *et al*., 2012; Cao *et al*., 2023). While RNAse-like effectors can also be found in other phytopathogenic fungi, this group of effectors appears to be strongly expanded within the *Blumeria* genus where it comprises more than half of the total effector complement (Müller *et al*., 2019; Cao *et al*., 2023; Seong & Krasileva, 2023). In this study, we found yet another three members of this effector superfamily to encode AvrPm60_1 and AvrPm60_2, thereby further highlighting the importance of the RNAse-like effectors in cereal powdery mildews. Interestingly, the previously described AvrPm3^a2/f2^ and the newly identified AvrPm60 effectors are part of the same effector family E008, whereas their corresponding NLRs belong to phylogenetically distinct NLR clades (Avni *et al*., 2022). The question remains, however, why RNAse-like effectors in general, and the E008 family specifically, are especially prone to be recognized by NLRs. It has been hypothesized that the high expression levels observed during early stages of infection for many RNAse-like *AVR*s, including *AvrPm3^a2/f2^* and the newly identified *AvrPm60* genes (Supplementary Figure S2), makes this class of effectors predestined to be recognized by the hosts immune system (Bourras *et al*., 2018; Müller *et al*., 2019).

Another characteristic observed with *AVRs* in *Bgt* is extensive copy number variation of *AVR* genes within the global pathogen population. For example, *AvrPm3^d3^* was found to occur in up to six copies in some isolates, while other *AVRs,* such as *AvrPm3^a2/f2^* or *AvrPm17,* were found to occur in one to four copies (Supplementary Figure S3) (Bourras *et al*., 2019; Müller *et al*., 2019; Müller *et al*., 2022). Interestingly, we found three neighboring, nearly identical *AvrPm60* genes in the *Pm60-*avirulent isolate CHVD_042201, with the majority of isolates within the global collection containing two or three copies (Figure 4a). We hypothesize that gain of virulence mutations affecting a single *AvrPm60* gene copy might therefore not suffice to overcome *Pm60* resistance. This is consistent with the observation that complete deletion of the *AvrPm60* locus, as observed in the *Pm60*-virulent isolates CHN_52_27, USA_2, USA_3 and USA_5, is a potent gain-of-virulence mechanism (Figure 4a,b). Interestingly, *AvrPm60* deletion is rare among Chinese *Bgt* isolates with only two out of a 63 analyzed isolates exhibiting complete absence of *AvrPm60* genes (Figure 4a, Table 1). This observation is in line with earlier findings showing that *Pm60* provides broad resistance against the Chinese *Bgt* population (Zou *et al*., 2018, Zou *et al*., 2022). By contrast, deletion of *AvrPm60* genes is relatively common among *Bgt* isolates from the US (Figure 4a,b, Table 1). This finding is intriguing, given that we are not aware of any documented use of *Pm60* in agricultural production in the US. Nevertheless, we hypothesize that the frequent deletion of *AvrPm60* in this *Bgt* subpopulation could be the result of previous use of *Pm60* or *Pm60-like* resistance genes in this region, knowingly or unknowingly. Alternatively, it could be the consequence of the use of yet another resistance gene that recognizes the same effector proteins. Irrespective of underlying reasons for the frequent absence of *AvrPm60s* in the US *Bgt* population, we conclude that the resistance provided by *Pm60* is locally ineffective and therefore will likely provide only partial protection against *Bgt* infection in this geographic region.

A study by Zou and colleagues (2022) showed that the resistance provided by *Pm60a* and *Pm60b* resembles the one mediated by *Pm60* and concluded that the three alleles might have overlapping, albeit not identical, recognition spectra. In their study the authors showed that in particular *Pm60a* only provides resistance against a subset of isolates recognized by *Pm60* and *Pm60b*, which might indicate weaker or partially divergent recognition activity of *Pm60a* as compared to the other two alleles found in *T. urartu* (Zou *et al*., 2018; Zou *et al*., 2022). This agrees with the findings in this study where we could show that AvrPm60_1 and AvrPm60_2 trigger HR immune responses in the presence of the Pm60, Pm60a and Pm60b NLRs albeit at varying levels (Figure 4d,e). For AvrPm60_2 we observed strong HR responses upon co-expression with Pm60 and Pm60b but a weaker response with Pm60a (Figure 4d), corroborating the initial observations of Zou and colleagues (Zou *et al*., 2018; Zou *et al*., 2022). Furthermore, co-expression of Pm60a and Pm60b with AvrPm60_1 resulted in a significantly weaker HR response as compared to Pm60 (Figure 4e). Our findings indicate that the deletion of two LRR repeats in Pm60a results in a lower sensitivity towards AvrPm60_1 and AvrPm60_2 whereas the duplication of the same two LRR repeats only influences AvrPm60_1 recognition. In conclusion, these observations indicate that Pm60 outperforms its allelic variants Pm60a and Pm60b in their ability to recognize AvrPm60_2 and in particular AvrPm60_1 effectors and should therefore be prioritized for wheat breeding.

Interestingly several allele pairs of the Pm3 allelic series were shown to recognize the same AVR protein with differing recognition strength. The alleles Pm3a (strong) and Pm3f (weak) were shown to recognize AvrPm3^a2/f2^, Pm3b (strong) and Pm3c (weak) recognize AvrPm3^b2/c2^ and finally, Pm3s (strong) and Pm3m (weak) a so far unknown AvrPm3^m/s^ (Stirnweis *et al*., 2014; Bourras *et al*., 2015; Bourras *et al*., 2019). In the case of Pm3 diversity the underlying amino acid polymorphisms defining strong and weak alleles were identified and reside within the NB-ARC domain of the NLR (Stirnweis *et al*., 2014). In contrast, the weaker Pm60a and Pm60b alleles are defined by the lack or duplication of two LRR repeats, respectively, compared to the stronger Pm60 allele. Thus, different categories of polymorphisms within NLR proteins can influence their strength and ability to recognize AVR proteins, highlighting the importance of studying NLR diversity in order to define the most potent variants for wheat protection.

Multiple studies identified additional *Pm60* diversity in wild emmer wheat (WEW, *Triticum dicoccoides*) and defined 11 haplotypes that differed from *Pm60* found in *T. urartu* predominantly by single amino acid polymorphisms affecting the CC, the NB-ARC domain as well as the LRR repeats (Li *et al*., 2021; Wu *et al*., 2021; Wu *et al*., 2022). While resistance activity of some *Pm60* alleles in WEW was verified, the ability of the remaining *Pm60* diversity to recognize *Bgt* remains unknown. The identification of *AvrPm60_1* and *AvrPm60_2* therefore provides the opportunity to study, and potentially validate additional *Pm60* alleles from WEW as well as investigate the consequences of polymorphisms found within *Pm60* on its ability to recognize the two known *AvrPm60’s* and, potentially, additional effector proteins.

Indeed, the identification and cloning of *Pm60* and *AvrPm60* genes in combination with investigations into their natural diversity opens new avenues of research and provides the opportunity to study the consequences of individual polymorphisms. These investigations will be crucial to estimate the value of *Pm60* for breeding and deployment and could, as exemplified by the high frequency of *AvrPm60* deletion observed in the US *Bgt* population, inform and guide regional deployment of resistance alleles. These considerations exemplify how the *R*/*AVR* atlas envisioned by Hafeez and colleagues could improve wheat resistance breeding in the future and how new technologies for rapid and cost-efficient *AVR* identification, such as the AD assay described in this study, can pave the way for pathogen-informed resistance gene deployment.

## Methods

### *Bgt* isolates, sexual crosses and selection of F1 populations

The *Bgt* isolate CHVD_042201 (mating type MAT 1-2) was collected from a powdery mildew infected wheat field in Begnins, canton of Vaud, Switzerland in spring 2022 and subsequently single spore isolated twice in order to ensure a genetically uniform culture. The *Bgt* isolate CHN_52_27 (MAT 1-1) has previously been described in (Zeng *et al*., 2014; Praz *et al*., 2017). All other *Bgt* isolates used in this study have previously been described in (Sotiropoulos *et al*., 2022). *Bgt* isolates were maintained clonally on the susceptible wheat cultivar ‘Kanzler’. For conidiospore production, infected leaf segments were placed on food grade agar plates (0.5%, PanReac AppliChem) supplied with 4.23mM benzimidazole and incubated at 20°C as described previously (Parlange *et al*., 2011). The sexual cross between CHVD_042201 and CHN_52_27 was performed as previously described (Parlange *et al*., 2015) by co-infecting the susceptible wheat cultivar ‘Kanzler’. Leaf segments harboring chasmothecia were harvested and dried at room temperature for several weeks. For ascospore ejection, dried chasmothecia were exposed to high humidity using Whatman filter paper soaked in sterilized water for up to 10 days and arising F1 progeny collected and grown on the susceptible wheat cultivar ‘Kanzler’. Resulting F1 conidiospores were subsequently subjected to bulk selection on wheat cultivars ‘Kanzler’ (no selection), ‘Asosan/8*CC’ (*Pm3a* selection) or ‘Kn199 + *Pm60’* transgenic plants (*Pm60* selection). Selected F1 conidiospore bulks were harvested after 10 days and fungal DNA extracted as described below.

### Plant material, virulence scoring

The *Pm3a* containing line ‘Asosan/8*CC’ has been previously described in (Bourras *et al*., 2015). The ‘Kn199 + *Pm60’* transgenic line, expressing *Pm60* under endogenous promoter and terminator sequences in the susceptible ‘Kn199’ background, has been previously described in (Zou *et al*., 2018).

To determine virulence phenotypes, leaf segments of the cultivars ‘Kanzler’, ‘Asosan/8*CC’ or ‘Kn199+*Pm60*’ were placed on agar plates as described above, infected with the indicated *Bgt* isolates and disease phenotypes imaged at 8-9 days post infection. Virulence scoring was performed on at least six biological replicates for each tested interaction. Representative images were chosen for the depiction of virulence phenotypes throughout the manuscript.

### DNA sequencing and genome assembly

Fungal DNA was extracted from conidiospores using a previously described CTAB/phenol-chloroform extraction procedure (Bourras *et al*., 2015). For the CHVD_042201 genome assembly, 5µg of high molecular weight DNA was used for library preparation and PacBio HiFi sequencing was performed on the PacBio Sequel Ile platform using a 30h movie at the Functional Genomics Center Zurich (FCGZ). Resulting PacBio HiFi raw data is available at the sequence read archive (SRA, accession: PRJNA1131794).

PacBio HiFi reads were assembled using HiCanu (Nurk *et al*., 2020), HiFlye (Kolmogorov *et al*., 2019) and hifiasm (Cheng *et al*., 2021) as described in Supplementary Note 1. HiCanu assembly was performed with the options genomeSize=141m -pacbio-hifi. HiFlye assembly was performed with the options -g 141m –pacbio-hifi. Hifiasm assembly was performed with the options -t16 -l0 -f0 --hg-size 141m. Subsampling of PacBio HiFi reads was performed with seqtk sample command (https://github.com/lh3/seqtk).

Whole genome alignments of assemblies against Bgt_genome_v3_16 was performed using the mummer suite (v4.0.0) (Marcais *et al*., 2018) using the command nucmer. Subsequent plots were rendered with the mummerplot command using the following specification: --filter --color –png. Subsequently, plots were produced with gnuplot.

Blast searches of PacBio HiFi reads against the mitochondrial sequence of CHE_96224 (Genebank: MT880591.1) were performed using the BLAST+ suite (v2.12.0) (Camacho *et al*., 2009) with the following specifications: - qcov_hsp_perc 50 and all reads that aligned to the mitochondrial genome were retained. Subsequently, these reads were used to perform an assembly using hifiasm with the -l0 option.

The final assembly was polished with Illumina reads from isolate CHVD_042201 that were mapped against the assembly with the method described in (Kunz *et al*., 2023). Subsequent polishing was performed with Pilon (v1.24) (Walker *et al*., 2014) with the following specifications: --fix bases –changes. The polished genome assembly of isolate CHVD_042201 is available as Bgt_CHVD_042201_genome_v1 on Zenodo (https://zenodo.org/records/11233413)

### Annotation

Annotation of the Bgt_CHVD_042201_genome_v1 was performed using the MAKER2 software (v2.31.11) (Cantarel *et al*., 2008), available from the European Galaxy server (https://usegalaxy.eu/). Repeat masking of the genome was achieved using the TREP database (trep-db_nr_Rel-19.fasta and trep-db_proteins_Rel-19.fasta) available at https://trep-db.uzh.ch/. We used the prot2genome option of MAKER2 to create a homology-based draft annotation of Bgt_CHVD042201_genome_v1 in two rounds. The first round was performed using the predicted proteome of CHE_96224 (v4_23, https://zenodo.org/records/7018501), and the second round using the proteome of *Bh* strain DH14 (https://github.com/lambros-f/blumeria_2017/tree/master/annotation_genome_dh14) . Genes predicted in the second round were only included if they did not overlap with any genes predicted in the first round. The annotation of of Bgt_CHVD042201_genome_v1 is available on Zenodo: (https://zenodo.org/records/11233413)

### Avirulence depletion assay

DNA from the *Bgt* isolates CHVD_042201 and CHN_52_27 was sequenced at the Functional Genomics Center Zurich (FGCZ). Sequencing libraries were generated using the Illumina Trueseq Nano protocol and sequencing was performed on the Illumina Novaseq 6000 platform. DNA from unselected, *Pm3a-* or *Pm60-*selected F1 bulks was sequenced to a coverage of ∼120X with our commercial partner Novogene UK on a NovaSeq X Plus platform. Mapping of Illumina reads was performed as described previously (Kunz *et al*., 2023). For the analysis of the bulk-sequencing data, we established a pipeline specifically tailored towards the haploid genomic structure of *Bgt*. All steps of the pipeline were executed using a custom Python script, available on Github: https://github.com/MarionCMueller/AD-assay.

In detail, the pipeline first identified high-quality single nucleotide polymorphisms (SNPs) in the two parental isolates, CHVD_042201 and CHN_52_27 as follows: Illumina mapping files were simultaneously used to perform SNP calling with FreeBayes (v1.3.6) (Garrison & Marth, 2012), using the following options: --haplotype-length 0, --min-alternate-count 20, --min-alternate-fraction 0, --pooled-continuous, and --limit-coverage 400. The resulting polymorphic sites were further filtered to retain only those SNP positions where both parents had at least 10 reads and exhibited opposite genotypes. Genotypes were only accepted if 95% of the reads supported the genotype. Subsequently, the pipeline identified SNPs in the alignment files (BAM files) of the unselected and selected bulk only at the polymorphic positions between the parental isolates using FreeBayes with the options: --haplotype-length 0, --min-alternate-count 1, --min-alternate-fraction 0, --pooled-continuous, and --report- monomorphic.

The subsequent statistical analysis was conducted in R using the functions provided in the BSA_Blumeria_functions.R object available on GitHub at https://github.com/MarionCMueller/AD-assay. To ensure the removal of sites exhibiting copy number variation in one of the parental isolates, the read coverage at each marker position was analysed. Positions with sequencing coverage either above or below twice the standard deviation of the coverage of all sites were excluded. Next, the G.test() function of the R package RVAideMemoire was utilized to assess the deviation of the parental SNP ratio from an expected 1:1 distribution for unselected marker positions. Finally, resulting p-values were averaged over 10 SNPs using the runner() command. Scripts used for analysis are available as an R Markdown object from Github (https://github.com/MarionCMueller/AvrPm60).

### Cloning

For expression of fungal effectors in *N. benthamiana*, the predicted signal peptide (SignalP4.0 (Petersen *et al*., 2011) was removed and replaced by a start codon. The remaining coding sequences of all effector candidates were codon-optimized for expression in *N. benthamiana* based on the codon-optimization tool of Integrated DNA technologies (https://eu.idtdna.com). Optimized sequences were gene synthesized with gateway compatible flanking attL sites with our commercial partners (BioCat GmbH https://www.biocat.com; Thermo Fisher Scientific https://www.thermofisher.com). The resulting gateway compatible entry clones were subsequentially mobilized into the binary expression vector pIPKb004 (Himmelbach *et al*., 2007) using Invitrogen LR clonase II according to the manufacturer.

For expression of *Pm3a* in *N. benthamiana* we made use of a pIPKb004-Pm3a-HA construct that has been previously described (Bourras *et al*., 2015; Bourras *et al*., 2019). For the expression of *Pm60*, *Pm60a* and *Pm60b* in *N. benthamiana* we first amplified the *Pm60* coding sequence from pEarlyGate-Pm60 described in (Zou *et al*., 2018) using KAPA HiFi Polymerase (KAPA Biosystems) with the primers listed in Supplementary Table 3 and cloned the resulting PCR amplicon into the gateway compatible entry vector pDONR221 using BP clonase (Invitrogen), resulting in pDONR221-Pm60. The polymorphic regions defining the *Pm60a* and *Pm60b* alleles were gene synthesized with our commercial partner Thermo Fisher Scientific (https://www.thermofisher.com) and introduced into the *Pm60* coding sequence using PCR amplification with KAPA HiFi Polymerase (KAPA Biosystems) and the primers listed in Supplementary Table 3 applying the In-Fusion cloning method (Takara Bio) according to the manufacturer, resulting in pDONR221-Pm60a and pDONR221-Pm60b. The entry clones of *Pm60*, *Pm60a* and *Pm60b* were mobilized into pIPKb004 as described above. The sequences of all DNA fragments produced by gene-synthesis can be found in Supplementary Table 4. All constructs in the binary expression vector pIPKb004 were transformed into *A. tumefaciens* strain GV3101 using freeze-thaw transformation (Weigel & Glazebrook, 2006).

### Co-expression of *AVR* candidates and *R* genes for HR quantification in *N. benthamiana*

*Agrobacterium*-mediated transient expression of effector candidates and resistance genes in *N. benthamiana* was achieved with the detailed protocol described in (Bourras *et al*., 2019). For co-expression of *AVR* candidates and corresponding *R* genes, *Agrobacteria* OD1.2 were mixed in a 4:1 ratio (AVR:R) prior to infiltration. HR imaging and quantification was performed 4-5 days after *Agrobacterium* infiltration using a Fusion FX imaging system (Vilber Lourmat https://www.vilber.com/) and the Fiji software (Schindelin *et al*., 2012) as described previously in (Bourras *et al*., 2019).

### Gene expression analysis

To quantify gene expression, we used previously published dataset of *Bgt* isolates CHE_96226, CHE_94202, GBR_JIW2, ISR_7 and CHN_17_40 (Praz *et al*., 2018; Müller *et al*., 2022; Kunz *et al*., 2023). Accession numbers of the RNAseq libraries are listed in Supplementary Table 5. RNAseq reads were pseudoaligned to the CHVD_042201 CDS using the salmon software v1.4.0 (Patro *et al*., 2017). First, CHVD_042201 CDS (Bgt_CHVD042201_CDS_v1_1.fasta) was indexed using the command salmon index. Then, single or paired end reads were quantified with the command salmon quant -l A. Subsequently, raw read counts were converted to RPKM values using the edgeR package 3.40.2 (Robinson *et al*., 2010) using the rpkm() command. Plots were generated with ggplot2 v3.4.3 using a custom R script in RStudio v2023.03.0+386 (Wickham, 2009; RStudio- Team, 2018). An R Markdown script detailing all code used in this analysis to conduct read count, quantification and plot generation is available on Github (https://github.com/MarionCMueller/AvrPm60).

### Copy number variation

Analysis of copy number variation was performed based on previously published *Bgt* diversity data available described in (Sotiropoulos *et al*., 2022; Kloppe *et al*., 2023). Sequencing reads were aligned to the *Bgt* reference genome Bgt_genome_v3_16 using the method described in (Sotiropoulos *et al*., 2022). Subsequently read coverage for each gene was extracted and normalised to the average coverage of all genes in the genome with a previously published script available on Github (https://gist.github.com/caldetas/24576da33d1ff91057ecabb1c5a3b6af). Genes exhibiting a normalized coverage below 0.1 were considered deleted. A table with normalized coverage values for all isolates is available here: (https://github.com/MarionCMueller/AvrPm60). Data was visualized and further analysed with a custom R script available on Github (https://github.com/MarionCMueller/AvrPm60).

### AlphaFold 3 Modelling

For AlphaFold 3 modelling of AVRPM3A2/f2 and AVRPM60, the signal peptides were removed based on predictions from SignalP 5.0 (https://services.healthtech.dtu.dk/services/SignalP-5.0/). The structures were then predicted using the AlphaFold 3 server (https://alphafoldserver.com/) (Abramson *et al*., 2024). Visualization of the structures was performed using ChimeraX software (Meng *et al*., 2023).

## Author contributions

L.K., M.C.M, and B.K. designed the research. L.K. and M.C.M wrote the manuscript. B.K., R.H. and F. M. edited the manuscript. L.K. and H.Z. performed the experiments. M.C.M., L.K., A.G.S. performed bioinformatic analyses.

L.K. and M.C.M. analysed data. J.J. and F.M. provided powdery mildew isolates. S.Z. and D.T. provided transgenic wheat lines. All authors read and approved the final manuscript.

## Data availability

PacBio HiFi reads for isolate CHVD_042201 are available from the sequence read archive (SRA, accession: <u>PRJNA1131794</u>). Genome assembly and annotation of isolate CHVD_042201 are available from Zenodo: https://zenodo.org/records/11233413. Illumina data for the two parental isolates used for the AD assay are available as follows: CHVD_042201 is available from ENA (accession: PRJEB75381) and CHN_52_27 from SRA (accession: SRR29761601). Illumina sequencing data from the bulks generated by the AD assay are available at SRA (accession: PRJNA1125842). Scripts used to analyze the bulk sequencing data generated by the AD assay are available from https://github.com/MarionCMueller/AD-assay.

