## Supplementary Material for "Avirulence depletion assay: combining *R* gene-mediated selection with bulk sequencing for rapid avirulence gene identification in wheat powdery mildew"

**Supplementary Note 1: PacBio HiFi-based telomere-to-telomere assembly of Swiss *Bgt* isolate CHVD_042201**

To create a *de novo* assembly of *Bgt* isolate CHVD_042201 we sequenced its genome to a coverage of approximately 100X using CCS HiFi reads on the PacBio Sequel II system. Using the resulting 12.8M PacBio CCS HiFi reads, we first benchmarked three commonly used genome assemblers specifically designed for HiFi reads: HiFlye (Kolmogorov *et al.*, 2019), HiCanu (Nurk *et al.*, 2020), and hifiasm (Cheng *et al.*, 2021). To assess the quality of the resulting assemblies, we performed whole genome alignments against the *Bgt* reference genome (Bgt_genome_v3_16, (Müller *et al.*, 2019)), which has assembled chromosomes that were resolved based on information of a genetic map and BAC-end sequences, and subsequently evaluated the ability of each assembler to resolve chromosome structures and complex regions such as centromeres. Notably, the use of the hifiasm assembler resulted in better resolution of both chromosomes and centromeres. Hifiasm successfully assembled seven out of the eleven chromosomes completely, compared to five for HiCanu and two for HiFlye. Additionally, all eleven centromeric regions were assembled without gaps in the hifiasm assembly, whereas HiCanu and HiFlye only assembled nine and six centromeres, respectively. Consequently, we chose to use the hifiasm assembler for all subsequent analyses.

Investigating the regions causing alignment breaks in the hifiasm assembly revealed that they corresponded to two rDNA clusters located on Chr-05 (5.8S rDNA) and Chr-09 (18S/28S rDNA), a tandem repeat cluster specific to *Blumeria* on Chr-04 (Müller *et al.*, 2019), and a candidate effector cluster on Chr-01. Strikingly, all these regions shared a common characteristic: due to their repetitive nature, they exhibited higher read coverage than the remaining regions of the genome. We hypothesized that subsampling of sequencing reads might improve the assembly resolution of these complex regions. Indeed, subsampling to approximately 30, 40, 50, and 60X coverage improved the assembly of the effector cluster on Chr-01 as well as the 5.8S rDNA cluster on Chr-05 for multiple subassemblies. For instance, the alignment break on Chr-01 was resolved in 9 out of the 20 subassemblies. Similarly, the rDNA cluster on Chr-05 was assembled in 4 out of 20 subassemblies. However, the tandem repeat cluster on Chr-04 and the rDNA cluster on Chr-09 were not resolved in any of the subassemblies. To create the final CHVD_042201 assembly, we utilized the initially produced hifiasm assembly based on all sequencing reads. Subsequently, we filled above-mentioned gaps with the corresponding sequences originating from one of the subsampled assemblies. For the effector cluster on Chr-01 all 9 subassemblies that resolved the cluster were identical, thus a random one was chosen to fill the gap. For the 5.8S rDNA cluster on Chr-05 the assembly with the longest repeat cluster was selected. This strategy allowed us to assemble nine of the eleven *Bgt* chromosome gapless (i.e. telomere-to-telomere), with Chr-09 and Chr-04 each containing a single remaining sequence gap corresponding to the 18S/28S rDNA cluster and the tandem repeat cluster, respectively.

Hifiasm initially also failed to assemble the mitochondrial genome, likely due to the fact that sequence coverage of the mitochondrion was exceeding the coverage of the nuclear genome. In the primary assembly, 275 contigs corresponded to the mitochondrial genome. We removed these contigs from the initial assembly, and identified all HiFi reads corresponding to the mitochondrion by aligning the reads to previously published mitochondrial genomes of isolates CHE_96224 using BLAST. Subsequently, we used hifiasm to separately assemble all sequencing reads corresponding to the mitochondrial genome, which resulted in a single circular contig of 102.7 kb, as well as 16 short contigs that were not incorporated into the final assembly. We also removed 157 contigs that contained additional sequences corresponding to rDNA clusters and five contigs that had BLAST hits to the genus *Penicillium*, likely indicating contamination. After one round of polishing the assembly using short-read DNA sequencing of isolate CHVD_042201 with the Pilon software (Walker *et al.*, 2014), the final assembly of CHVD_042201 consisted of 142.9 Mb, resolved into eleven *Bgt* chromosomes, a 100 kb mitochondrial genome, and five short unassembled contigs. We found 22 occurrences of the telomere repeat sequence at the ends of the eleven chromosomes of CHVD_042201.

Alignment of short-read DNA sequences of CHVD_042201 against the final genome assembly, followed by read coverage analysis in 500 bp windows, indicated that only 446 kb of the assembly sequence might still contain collapsed sequences. This is a significant improvement compared to the 2.24 Mb of collapsed sequence in the *Bgt* reference genome Bgt_genome_v3_16 of isolate CHE_96224 (Müller *et al.*, 2019). In the CHVD_042201 assembly, the collapsed regions are mostly confined to the highly repetitive tandem duplicated rDNA cluster and the tandem repeat cluster on Chr-04. We estimated this collapsed region to account for an additional 5.6 Mb of sequences, bringing the total size of the CHVD_042201 genome to an estimated 147.9 Mb (Supplementary Table S2).

Finally, we used the maker2 software to create a homology-based draft annotation for CHVD_042201. We used the predicted proteomes of both *Bgt* reference genome CHE_96224 and *Blumeria hordei* reference genome DH14 (Frantzeskakis *et al.*, 2018) to predict gene models in CHVD_042201. This strategy resulted in a draft genome annotation of 9’932 genes in the Bgt_CHVD042201_genome_v1.

The genome assembly and annotation of CHVD_042201 are available from Zenodo: https://zenodo.org/records/11233413

### Supplementary Note 2: Performing the AD assay based on alternative reference genomes

In this study, we developed a bioinformatics pipeline to analyze data generated by the AD assay and identify regions showing deviation from a 1:1 parental genotype ratio in experimentally generated bulk sequencing data in the haploid organism *Bgt*. The experimental setup in this study allowed us to perform our analysis based on the novel high-quality reference genome of CHVD_042201, which represented the avirulent parental isolate on *Pm3a* and *Pm60* resistance genes under investigation. However, this particular experimental setup might not always be achievable due to the significant cost and time investment required to create a high-quality genome assembly of *Bgt.*

For *Bgt*, four high-quality genome assemblies are publicly available, including the CHVD_042201 assembly presented in this study (Müller *et al.*, 2019; Müller *et al.*, 2021; Müller *et al.*, 2022). Our goal was to develop the pipeline for the AD assay to run on any *Bgt* reference genome, in order to maximize the success of *AVR* gene identification even without a high-quality reference genome of the avirulent isolate used for the genetic cross. Therefore, we tested our pipeline using the *Bgt* reference genome assembly of isolate CHE_96224 (Bgt_genome_v3_16) as the basis of the AD assay and compared the results with the initial analysis relying on the CHVD_042201 genome assembly.

In a first step, we mapped the Illumina sequencing data of both parental isolates (CHVD_042201, CHN_52_27) and the selected and unselected bulks against the reference assembly Bgt_genome_v3_16. Similar to our initial analysis using the CHVD_042201 genome, we then proceeded to call SNPs between the two parental isolates CHVD_042201 and CHN_52_27 in order to use them as genetic markers for the analysis of the bulks. Based on this strategy, we identified 193'871 SNP markers segregating between CHVD_042201 and CHN_52_27 when using the Bgt_genome_v3_16, compared to 198'027 SNP markers identified when using the Bgt_CHVD042201_genome_v1 as a reference, showing that the number of markers identified with both approaches is comparable.

The subsequent genotype depletion analysis, using the G-test to identify deviations from the parental genotype ratio of 1:1, was consistent across the analyses with either of the two reference assemblies. Both approaches identified a single region on *Bgt* Chr-06 for the *Pm3a*- and *Pm60*-selected bulks, which was absent in the non-selected bulk created on the susceptible cultivar 'Kanzler' (Supplementary Figure 1, Supplementary Table 2). This demonstrates that the AD pipeline is effective on an alternative reference genome assembly that does not represent the avirulent parental isolate used for the genetic cross and is therefore broadly applicable.

We analyzed the regions showing the strongest depletion signal (i.e. 95% and 90% of reads originating from the virulent parent) from both analyses. For *AvrPm3*$^{a2/f2}$, the interval based on Bgt_genome_v3_16 was 274 kb, significantly larger than the 25 kb interval identified using the CHVD_042201 assembly. In contrast, the interval for *AvrPm60* was 535 kb based on Bgt_genome_v3_16 compared to the 667 kb interval identified in our initial analysis based on the CHVD_042201 assembly. In both cases, the *AvrPm3*$^{a2/f2}$ and *AvrPm60* genes were located within the identified genomic regions, thus showing that the AD assay identified the correct genomic location.

In summary, we demonstrate that the AD assay successfully identifies the *AvrPm3*$^{a2/f2}$ and *AvrPm60* loci when Bgt_genome_v3_16 is used as the reference genome, suggesting that the AD pipeline can readily be used with other *Bgt* reference genomes.  However, we recommend using a reference genome from an avirulent isolate on the *R* gene under investigation in order to ensure efficient avirulence candidate gene definition. This is best exemplified by the case of the *AvrPm60* genes, where the identified virulence allele in isolate CHN_52_27 is defined by a large-scale deletion encompassing of all three *AvrPm60* copies (Figure 3c). In such a scenario, the use of a reference genome from a virulent strain carrying the same deletion would fail to identify *AvrPm60* candidate genes.

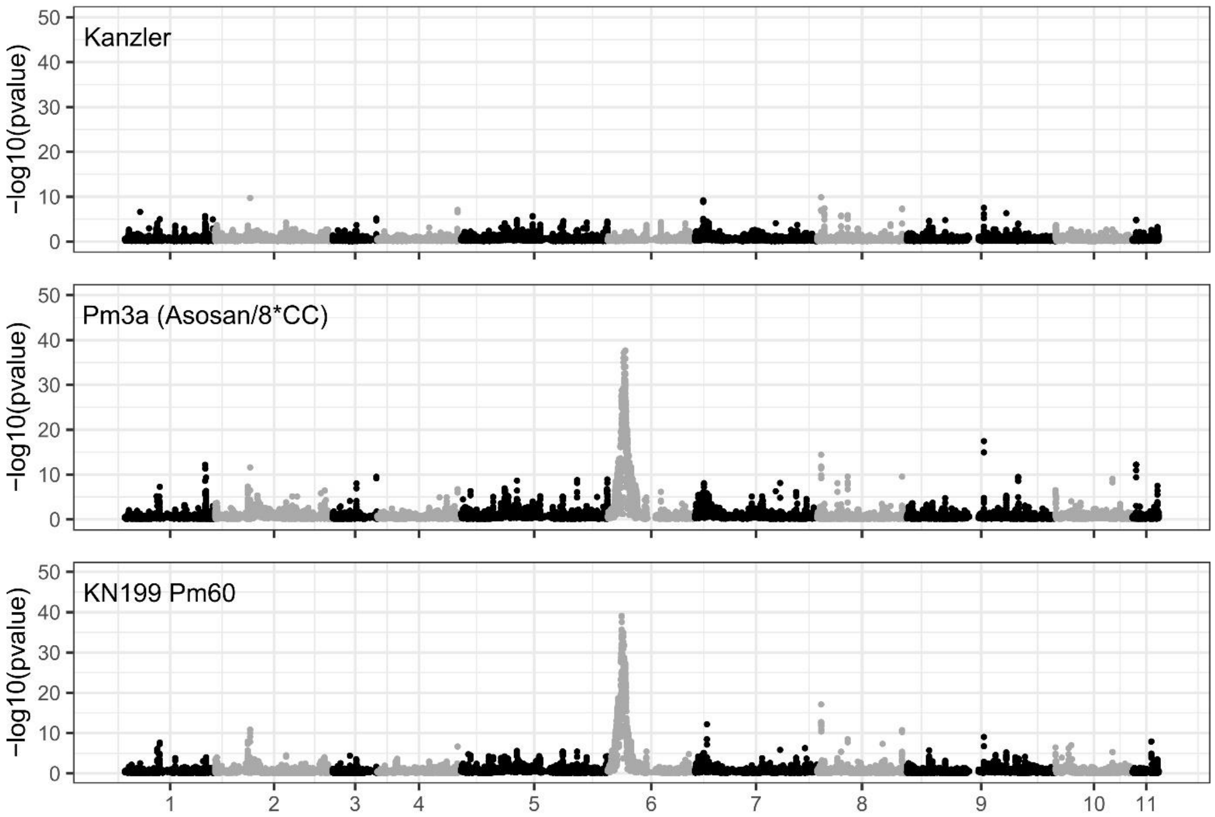


**Supplementary Figure S1: The AD assay performed on reference genome Bgt_genome_v3_16.**
The AD assay based on the CHVD042201 x CHN5227 cross was performed using the reference genome of Bgt_genome_v3_16 (CHE_96224). Cultivars used for selection are indicated in the respective plots. The Y-axis indicates –log10(p-values) of G-test statistics used to test for a deviation from 1:1 parental genotype ratio expected in the absence of selection. G-test values were averaged over 10 neighbouring SNPs.

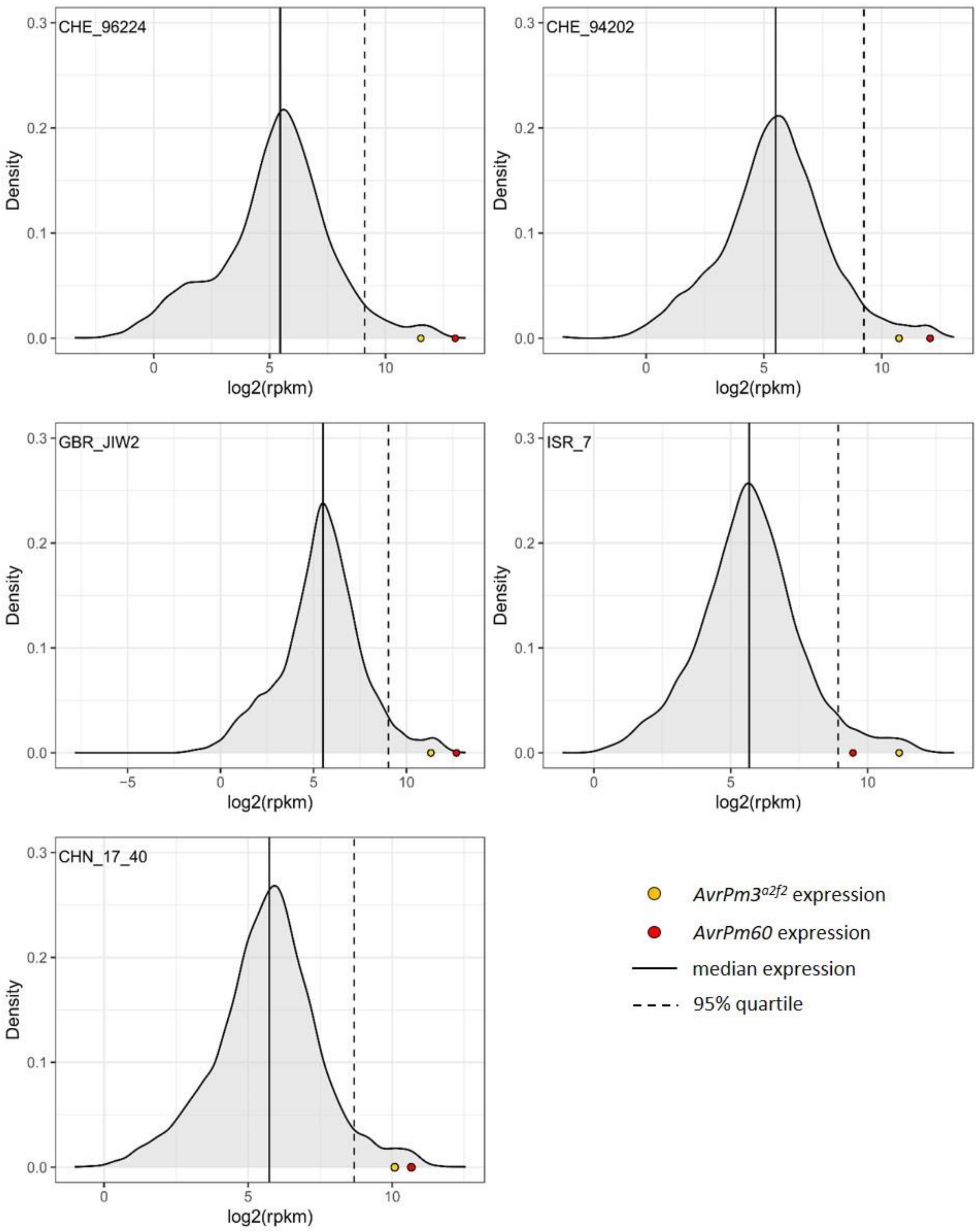


**Supplementary Figure S2: *$AvrPm3^{a2/f2}$* and *AvrPm60* are among the highest expressed genes in five *Bgt* isolates.** Distribution of all expressed (rpkm >0) genes in the *Bgt* isolates CHE_96224, CHE_94202, GBR_JIW2, ISR_7 and CHN_17_40 are displayed as log2(rpkm) values. The coding sequence of isolate CHVD_042201 was used as a reference for quantification. All RNA sequencing data were obtained at 2 days post infection on the susceptible wheat cultivar 'Chinese Spring'. The solid line and the dashed line indicate the median and the 95% quartile of all expressed genes, respectively. Expression of *$AvrPm3^{a2/f2}$ (CHVD042201-04754)* and *AvrPm60* are indicated by a yellow and red dot respectively. *AvrPm60* expression represents the sum of the rpkm values of all three *AvrPm60* genes in the reference isolate CHVD_042201 (*CHVD042201-04743*, *CHVD042201-04745* and *CHVD042201-04747).*

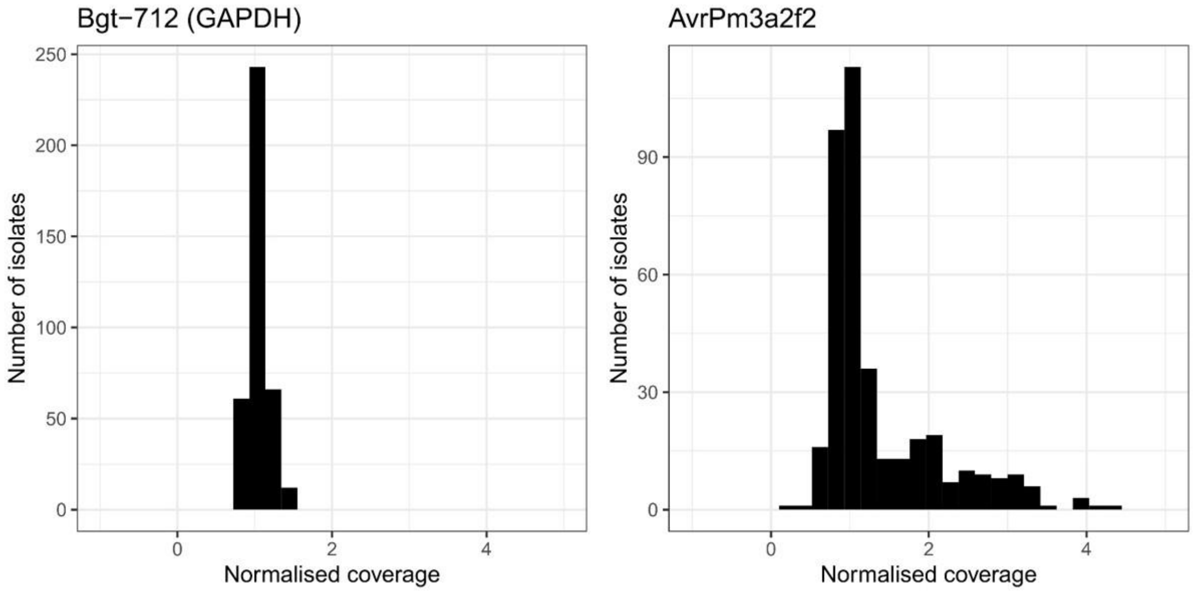


**Supplementary Figure S3: Distribution of normalized genomic coverage of the *GAPDH* and *AvrPm3$^{a2/f2}$* genes.** For each gene, the genomic coverage of sequencing reads was normalised to the coverage of all genes in the genome.

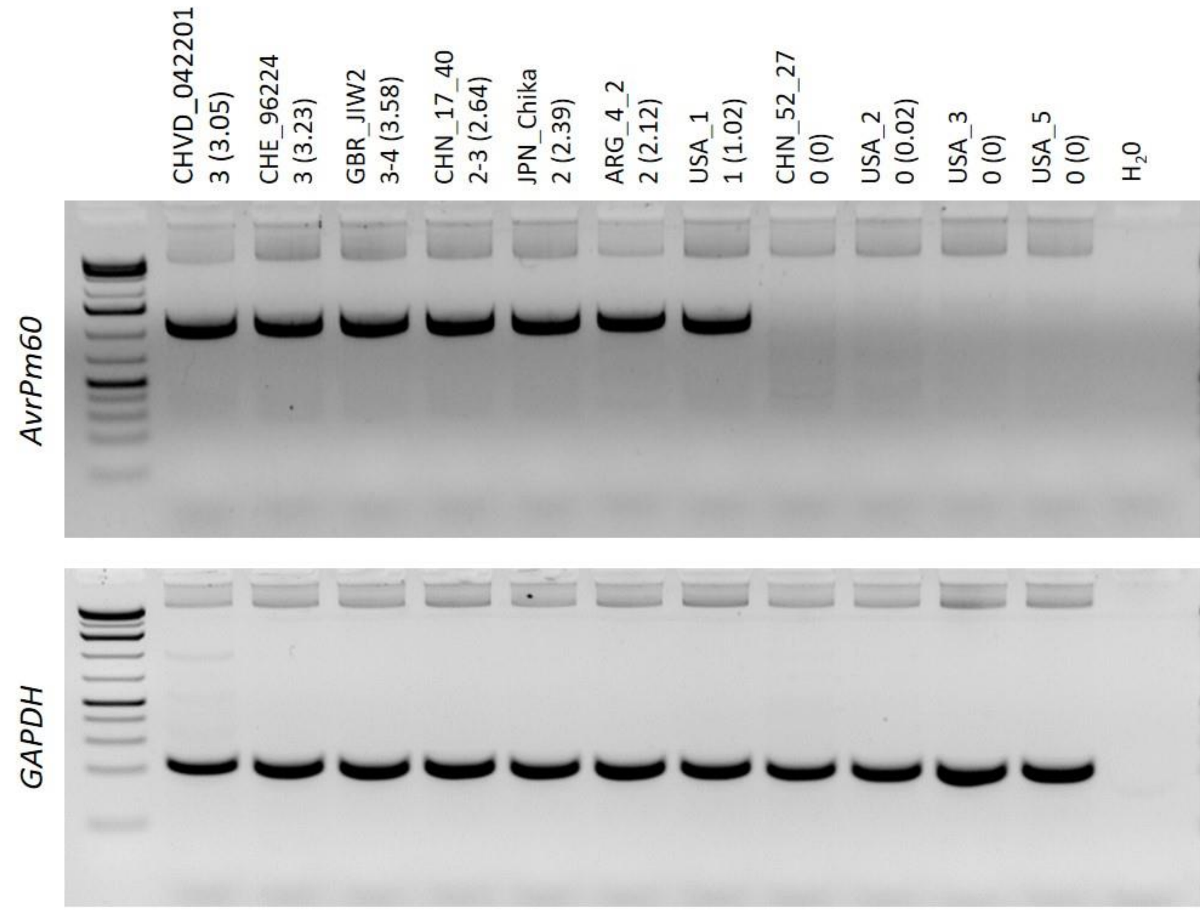


**Supplementary Figure S4: PCR verification of the *AvrPm60* gene deletion in *Bgt* isolates originating from China and the US.** *AvrPm60* specific primers were designed in conserved regions flanking the three *AvrPm60* genes (CHVD042201-04743, CHVD042201-04745 and CHVD042201-04747) in CHVD_042201 with a predicted amplicon size of 1179bp. Fungal *GAPDH* served as a positive control. The estimated number of *AvrPm60* gene copies according to sequencing coverage analysis are indicated next to the isolate name with normalized sequencing coverage indicated in brackets. The isolates CHVD_042201 and CHN_52_27 used for initial *AvrPm60* identification are shown as comparison.

**Supplementary Table 1: Genome stastistics of Bgt_CHVD042201_genome_v1**

| *Bgt* isolate | CHVD_042201 | CHE_96224 |
|---|---|---|
| Assembly | Bgt_CHVD_042201_genome_v1 | Bgt_genome_v3_16[a] |
| Assembly size | 142'913'105 | 140'575'254 |
| Largest scaffold | 16'394'915 | 4'473'759 |
| Number of sequence gaps | 2 | 357 |
| Chromosome number | 11 | 11 |
| Telomeric repeats | 22 | 7 |
| Number of genes | 9’932 | 8’581 |

[a] see (Müller *et al.*, 2019)

**Supplementary Table 2: Results of the AD assay using different reference genomes**

| Reference genome | Cross | Selection line | Selection strength | Start | End | Interval size |
|---|---|---|---|---|---|---|
| Bgt_CHVD042201_genome_v1 | CHVD_042201 x CHN_52_27 | Asosan/8*CC | 90% | 1911738 | 2647215 | 735477 |
| Bgt_CHVD042201_genome_v1 | CHVD_042201 x CHN_52_27 | Asosan/8*CC | 95% | 2272209 | 2297683 | 25474 |
| Bgt_CHVD042201_genome_v1 | CHVD_042201 x CHN_52_27 | Kn199 Pm60 | 90% | 1765214 | 2701804 | 936590 |
| Bgt_CHVD042201_genome_v1 | CHVD_042201 x CHN_52_27 | Kn199 Pm60 | 95% | 1904112 | 2570698 | 666586 |
| Bgt_genome_v3_16 | CHVD_042201 x CHN_52_27 | Asosan/8*CC | 90% | 1842969 | 2463023 | 620054 |
| Bgt_genome_v3_16 | CHVD_042201 x CHN_52_27 | Asosan/8*CC | 95% | 2101179 | 2375031 | 273852 |
| Bgt_genome_v3_16 | CHVD_042201 x CHN_52_27 | Kn199 Pm60 | 90% | 1369058 | 2483455 | 1114397 |
| Bgt_genome_v3_16 | CHVD_042201 x CHN_52_27 | Kn199 Pm60 | 95% | 1839679 | 2375034 | 535355 |

**Supplementary Table 3: List of primers used in this study.**

| Name | Sequence (5'-3') | Description/Purpose |
|---|---|---|
| HZ108 | GGGGACAAGTTTGTACAAAAAAGCAGGCTTC ATGGAATCGGCGATTGGCGCG | BP cloning to generate pDONR221-Pm60 |
| HZ109 | GGGGACCACTTTGTACAAGAAAGCTGGGTCC TACTCAAGTTCAAGTATCACATT | BP cloning to generate pDONR221-Pm60 |
| LK1102 | GAATGTGGCAACTTCTTTCTG | In-Fusion cloning to generate pDONR221-Pm60a |
| LK1103 | GGAAAAGCTTGATGGTGTG | In-Fusion cloning to generate pDONR221-Pm60a |
| LK1075 | ATCAGTTACAGTAAAAGAATGTGG | In-Fusion cloning to generate pDONR221-Pm60b |
| LK1076 | CAGAGCCATTGACTGCATG | In-Fusion cloning to generate pDONR221-Pm60b |
| LK1116 | GGATGTTTGGATCGCTC | *AvrPm60* specific primer |
| LK1117 | GTATATAGCAAAGGCTTGATG | *AvrPm60* specific primer |
| GAPDH_F | TGTCTTCCGAAACGCTGCTC | GAPDH specific primer (Bourras *et al.*, 2015) |
| GAPDH_R | AGTCCGTCCTCGACTGCTTGT | GAPDH specific primer (Bourras *et al.*, 2015) |

**Supplementary Table 4: Sequences of DNA fragments produced by gene synthesis.**

| Name | Sequence (5'-3') |
|---|---|
| CHVD042201-04754 (AvrPm3$^{a2/f2}$-A) | ATGGGTCCTGTCGCAAATGCTAGTTCTTATAAATGTCACGATAGAGTCATTGG TCCAGTGACTCTGAATGACCAGATCGAGAAGGCTTACCGTGAGGCCCTGGAG GCCGGCACGTCACCAAATGGACTTAGGAAAGGCCAGAAATTCGGTTCTCGTTA TTTCAACGTGATTCTTAAGAGGGGTGAGGAGAACATTAAAGTTGAATTCTTTG TTGGTATTAACTATCTCAAAGAGATTATTTACCTTCAGGCATATGTCCAGAGCG TTTTACTTGACTGTTACCCCACGACCGAGCGACCACAGTTGAACATTATCTTGC ACTAA |
| CHVD042201-04754_CHN52-27 (H36Q_G84E_E121D) (AvrPm3$^{a2/f2}$-B) | ATGGGTCCTGTCGCAAATGCTAGTTCTTATAAATGTCAAGATAGAGTCATTGG TCCAGTGACTCTGAATGACCAGATCGAGAAGGCTTACCGTGAGGCCCTGGAG GCCGGCACGTCACCAAATGGACTTAGGAAAGGCCAGAAATTCGGTTCTCGTTA TTTCAACGTGATTCTTAAGAGGGAAGAGGAGAACATTAAAGTTGAATTCTTTG TTGGTATTAACTATCTCAAAGAGATTATTTACCTTCAGGCATATGTCCAGAGCG TTTTACTTGACTGTTACCCCACGACCGATCGACCACAGTTGAACATTATCTTGC ACTAA |
| CHVD042201-04766 | ATGGACATTTCCAACTACTTGTGTGACCACGTCGTCTTGGACGCAAAGGACAT AGAGGCCGGAGTTGATAGAGCCTTTAGAACTAAAATGCAAGAGACGTTGGGG GCCTACGCCCCTGACGATTTCTATAACGAGGGCTCATATATCGTTAAGTATAA GTCTCCAAGGGTTAATATGGATGTCACTATTAAAATTGGTATCACGTTTAGTGA AGACGTACTTTACGTCAAAGCCGCAGGTGATGGGCAGGAGATAGATTGTCAC CCGACCGACAAACCGGCCACGACGAAACGTATCGTCCCGTAA |
| CHVD042201-04766_CHN52-27 (Y73N) | ATGGACATTTCCAACTACTTGTGTGACCACGTCGTCTTGGACGCAAAGGACAT AGAGGCCGGAGTTGATAGAGCCTTTAGAACTAAAATGCAAGAGACGTTGGGG GCCTACGCCCCTGACGATTTCTATAACGAGGGCTCATATATCGTTAAGAATAA GTCTCCAAGGGTTAATATGGATGTCACTATTAAAATTGGTATCACGTTTAGTGA AGACGTACTTTACGTCAAAGCCGCAGGTGATGGGCAGGAGATAGATTGTCAC CCGACCGACAAACCGGCCACGACGAAACGTATCGTCCCGTAA |
| CHVD042201-04767 | ATGGAGTTTTCTAACTATTTATGCGACCATATCGTAATAAATAAGAAAGACATA GAGTATTCAGTTGACCATGCTTTCAAAAAGCGTATGCAGGCTAACTTAGGAAA GTTCAAACCCGACGAAAAGTTCGGGACTGCTGGTTATATCACGAAATACAAGA CGGAGAAGGAAATTTTCGACGTTCATATTATTATCGAATATACAATCAACGAA GAAGTGATATCTGTTATCGCCAAAGGGCGTGGGCAGCAGGTGGTTTGCCACC CCACAGATCAGCCGGCCACGGAATATACAGAAGTAAGCGGGTCAGGCTAA |
| CHVD042201-04767_CHN52-27 (K74STOP) | ATGGAGTTTTCTAACTATTTATGCGACCATATCGTAATAAATAAGAAAGACATA GAGTATTCAGTTGACCATGCTTTCAAAAAGCGTATGCAGGCTAACTTAGGAAA GTTCAAACCCGACGAAAAGTTCGGGACTGCTGGTTATATCACGAAATACTAA |
| CHVD042201-04752 | ATGTATTCTCCAGTCCCTGTCGCCGAAGCCAGCAGTTATAAATGCCAAGATCG TGTAATTGGTCCTGTGACCCTTAACGATCAGATCAATAAGGCATACGCCGAGG CACAGAGTAATCAATCACGTGGACTGACCCGAGAGCAAATCTTTGCTTCCCGT CAATTTAGGGTAATCTTGACGAGAGACGGCGAGCGTATCTTAATCCAGTTTTA CCTGAGTATAAACAACGTCAAAGAGATTTTGTCCTTGCAAGCATACGTTATGA ACCAATTATTCACTTGTCATCCCACGATTGAACCACCACAATTAAACGAGGTTT TGCACTAA |

| | |
|---|---|
| CHVD042201-04752_CHN52-27<br>(I120T) | ATGTATTCTCCAGTCCCTGTCGCCGAAGCCAGCAGTTATAAATGCCAAGATCG<br>TGTAATTGGTCCTGTGACCCTTAACGATCAGATCAATAAGGCATACGCCGAGG<br>CACAGAGTAATCAATCACGTGGACTGACCCGAGAGCAAATCTTTGCTTCCCGT<br>CAATTTAGGGTAATCTTGACGAGAGACGGCGAGCGTATCTTAATCCAGTTTTA<br>CCTGAGTATAAACAACGTCAAAGAGATTTTGTCCTTGCAAGCATACGTTATGA<br>ACCAATTATTCACTTGTCATCCCACGACCGAACCACCACAATTAAACGAGGTTT<br>TGCACTAA |
| CHVD042201-04743;<br>CHVD042201-04745<br>(AvrPm60_1) | ATGGAAGGTAATTGCAATTACAAATGCGGTCCAGCAGTTATTGATGGTGATTA<br>TGTCAGGGAATGTGTCAAGTCTTACTACGAATATAAGATGCGTACCATAGATA<br>GGGATTACGGTCCCAACGATCATTTCACCACCGTCACCTTTCCGTTACAATACT<br>TGCACAAGGAGGAGATCATAACCGTACAAGTCAGCGCCGATTTTACGGCACTT<br>AGGGAAATAACCTCAGTTCGAGCAAGTGCTCTGGAGCAAGAGATAGAGTGTC<br>TGCCGACCCAATTGAAGCCTTCCTACGGAAAAGCCACCTGA |
| CHVD042201-04747<br>(E103G)<br>(AvrPm60_2) | ATGGAAGGTAATTGCAATTACAAATGCGGTCCAGCAGTTATTGATGGTGATTA<br>TGTCAGGGAATGTGTCAAGTCTTACTACGAATATAAGATGCGTACCATAGATA<br>GGGATTACGGTCCCAACGATCATTTCACCACCGTCACCTTTCCGTTACAATACT<br>TGCACAAGGAGGAGATCATAACCGTACAAGTCAGCGCCGATTTTACGGCACTT<br>AGGGAAATAACCTCAGTTCGAGCAAGTGCTCTGGGTCAAGAGATAGAGTGTC<br>TGCCGACCCAATTGAAGCCTTCCTACGGAAAAGCCACCTGA |
| Pm60a_fragment | CACACCATCAAGCTTTTCCCTGCTTCCCTCGAGACACTTGAGATTGAAGGAGA<br>GTCAGGCATGCAGTCAATGGCTCTGCTCAGCAATCTGAAATCCCTAAGGAGAC<br>TAGATGTCAGAAGATGCAGCATCACGTGCCATGGACTGCAGGACCTCGCGTG<br>CCTCCAATCAGTTACAGTAAAAGAATGTGGCAACTTCTTTCTG |
| Pm60b_fragment | CATGCAGTCAATGGCTCTGCTCAGCAATCTGAAATCCCTAAGGAGACTAGATG<br>TCAGAAGATGCAGCATCACGTGCCATGGACTGCAGGACCTCGCATGCCTCCAA<br>TCACTTACAGTACAAGACTGTGGCAACTTCTTTCCATGGCCTACCGAAGCAGCT<br>CACACCGTCAATCCTTTCCCTCACACCATCAAGCCTTTCCCTGCTTCCCTCGAGA<br>CACTTGAGATTGAAGGAGAGTTAGGCATGCAGCCAGTGGCTTTGCTCAGCAA<br>TCTGAAATCCCTAAGAAGACTAGATGTCAGAAGATGCAGCATCACGTGCCATG<br>GACTGCAGGACCTCGCATGCCTCCAATCACTTACAGTACAAGACTGTGGCAAC<br>TTCTTTCCATGGCCTACCGAAGCAGCTCACACCGTCAATCCTTTCCCTCACACCA<br>TCAAGCCTTTCCCTGCTTCCCTCGAGACACTTGAGATTGAAGGAGAGTTAGGC<br>ATGCAGCCAGTGGCTTTGCTCAGCAATCTGAAATCCCTAAGAAGACTAGATGT<br>CAGAAGATGCAGCATCACGTGCCATGGACTGCAGGACCTCGCGTGCCTCCAAT<br>CAGTTACAGTAAAAGAATGTG |

**Supplementary Table 5: RNA sequencing datasets used in this study.**

| Isolate | Type | Accession number | Publication |
|---|---|---|---|
| CHE_96224 rep1 | RNAseq | SRX3503528 | (Praz *et al.*, 2018) |
| CHE_96224 rep2 | RNAseq | SRX3503529 | (Praz *et al.*, 2018) |
| CHE_96224 rep3 | RNAseq | SRX3503530 | (Praz *et al.*, 2018) |
| CHE_94202 rep1 | RNAseq | SRX3503531 | (Praz *et al.*, 2018) |
| CHE_94202 rep2 | RNAseq | SRX3503532 | (Praz *et al.*, 2018) |
| CHE_94202 rep3 | RNAseq | SRX3503533 | (Praz *et al.*, 2018) |
| GBR_JIW2 rep1 | RNAseq | SRX3503534 | (Praz *et al.*, 2018) |
| GBR_JIW2 rep2 | RNAseq | SRX3503535 | (Praz *et al.*, 2018) |
| GBR_JIW2 rep3 | RNAseq | SRX3503536 | (Praz *et al.*, 2018) |
| ISR_7 rep1 | RNAseq | SRX17115170 | (Müller *et al.*, 2022) |
| ISR_7 rep2 | RNAseq | SRX17115171 | (Müller *et al.*, 2022) |
| ISR_7 rep3 | RNAseq | SRX17115172 | (Müller *et al.*, 2022) |
| CHN_17_40 rep1 | RNAseq | SRX18362804 | (Kunz *et al.*, 2023) |
| CHN_17_40 rep2 | RNAseq | SRX18362805 | (Kunz *et al.*, 2023) |
| CHN_17_40 rep3 | RNAseq | SRX18362806 | (Kunz *et al.*, 2023) |
